# Nobiletin Affects Circadian Rhythms and Oncogenic Characteristics in a Cell-Dependent Manner

**DOI:** 10.1101/2020.01.14.906420

**Authors:** Sujeewa S. Lellupitiyage Don, Kelly L. Robertson, Hui-Hsien Lin, Caroline Labriola, Mary E. Harrington, Stephanie R. Taylor, Michelle E. Farkas

## Abstract

The natural product nobiletin is a small molecule, widely studied with regard to its therapeutic effects, including in models of cancer. Recently, nobiletin has also been shown to affect circadian rhythms via their enhancement, resulting in protection against metabolic syndrome. We hypothesized that nobiletin’s anti-oncogenic effects are correspondingly accompanied by modulation of circadian rhythms. Concurrently, we wished to determine whether the circadian and anti-oncogenic effects of nobiletin differed across cell culture models of cancer. In this study, we assessed nobiletin’s circadian and therapeutic characteristics to ascertain whether these effects depend on cell line, which here also vary in terms of baseline circadian rhythmicity. Three cell culture models where nobiletin’s anti-cancer effects have been studied previously were evaluated here: U2OS (bone osteosarcoma), which possesses robust circadian rhythms; MCF7 (breast adenocarcinoma), which has weak circadian rhythms; and MDA-MB-231 (breast adenocarcinoma), which is arrhythmic. We found that both circadian and anti-cancer effects following nobiletin treatment were subtle in the U2OS and MCF7 cells. On the other hand, changes were clear in MDA-MB-231s, where nobiletin rescued rhythmicity, and substantially reduced oncogenic features. Based on these results and those previously described, we posit that a positive correlation exists between nobiletin’s circadian and therapeutic effects.

## Introduction

Nobiletin is a polymethoxylated flavone present in the peels of citrus fruits.^1^ It has been reported to yield therapeutic effects in a variety of disorders, including neurological, inflammatory, cardiac, and metabolic diseases, in addition to cancers. Recently, it has also been shown to affect circadian oscillations.^2,3^ We are interested in concomitant studies of nobiletin’s therapeutic and circadian effects. Previous studies have shown that nobiletin can improve memory in Alzheimer’s^4^ and motor deficits in Parkinson’s disease models,^5^ and can result in anti-depressant-like effects.^6^ It can also ameliorate adiposity, hyperlipidemia, hyperglycemia, and insulin resistance,^7^ attenuate lipid accumulation,^8^ protect against metabolic syndrome,^2^ reduce excessive inflammatory responses and restore epithelial barrier function in colitis,^9^ and protect against acute pancreatitis.^10^ For heart diseases, nobiletin results in neuroprotection of cerebral ischemia-reperfusion,^11^ lowers serum LDL/VLDL cholesterol,^12^ and has been shown to ameliorate cardiac dysfunction.^13^ Toward anti-aging, in mouse models, nobiletin has been shown to result in inhibition of bone resorption and maintenance of bone mass.^14^ On these accounts, nobiletin has the potential to serve as a therapeutic entity against numerous conditions.

Treatment with nobiletin has resulted in anti-oncogenic effects in a variety of cancer models by affecting major pathways including AKT, MAPK, TGF-β1/SMAD3, ERK, and JNK. In studies where it was shown to affect the AKT pathway, nobiletin was found to decrease tumor viability, weight, and volume in A2780/CP70 ovarian cancer xenograft models.^15^ It also reduced cell adhesion, invasion, and migration in the highly metastatic AGS gastric adenocarcinoma cell line,^16^ suppressed viability of PC-3 and DU-145 prostate cancer cell lines,^17^ and inhibited ACHN and Caki-2 renal carcinoma cell proliferation by cell cycle arrest in G0/G1 phase.^18^ Also, nobiletin causes anti-leukemic effects in HL-60 human acute myeloid leukemia cells,^19^ reduction of proliferation and migration in U87 glioma cells,^20^ and decreases cell proliferation in the low tumor-grade MCF7 cell line via G1 cell cycle block.^21^ It may induce apoptosis and inhibit cell migration via MAPK and/or MAPK/AKT related pathways.^22,23^ Similar anti-proliferative effects have been observed in high tumor grade MDA-MB-231^23^ and MDA-MB-468 human breast cancer cells following treatment with nobiletin.^24^ Nobiletin has also been shown to inhibit growth of metastatic nodules in the lungs of mice via the TGF-β1/SMAD3 pathway^25^ and decrease cell migration, angiogenesis, tumor formation, and progression in the bone osteosarcoma cell line U2OS via ERK and JNK pathways.^26^ Taken together, nobiletin is effective against different cancer types and exerts its effects via various pathways.

While identification of small molecules that can modulate circadian rhythms has garnered significant interest,^27^ it has recently been shown that many existing therapeutic compounds also affect circadian rhythms.^28^ In a similar fashion, nobiletin, which has been known for decades to elicit various beneficial health effects, has only recently been shown to alter circadian rhythms.^2,3^ Circadian rhythms are 24-hour rhythmic activity cycles. At the molecular level, they are regulated by CLOCK, BMAL, PER, and CRY proteins, which exist in a negative feed-back loop.^29^ CLOCK and BMAL are positive components that result in gene expression, including of PER and CRY, which act as negative regulators. Nobiletin has been shown to activate a critical component of a secondary loop, Retinoid Acid-related Orphan Receptor (ROR),^2^ which binds to the *Bmal1* promoter, activating its transcription.^29^ While other targets may exist, and at least two have been computationally predicted,^23,30^ they have yet to be determined. Two prior studies have assessed nobiletin’s circadian effects. In the first, mouse embryonic fibroblasts (MEFs) from heterozygous *Clock* knockout mice and mouse ear fibroblasts from *PER2∷LucSV* reporter mice showed increased *PER2:Luc* amplitudes, which were confirmed to be the result of increased ROR activity.^2^ In a second study, nobiletin similarly increased amplitudes but also lengthened the periods of *PER2::Luc* in MEFs, while inducing phase delays in liver slices from *PER2::Luc* knock-in mice.^3^ However, despite nobiletin’s therapeutic applications, to our knowledge, its simultaneous effects on circadian rhythms have been addressed only in a single study.^2^ Therefore, here we assess nobiletin’s circadian and therapeutic effects in parallel, under the same treatment conditions and models.

Given the substantial evidence of nobiletin’s anti-cancer efficacy, we considered it important to evaluate its circadian effects these systems. In this study, we used three cell culture cancer models, each of which nobiletin has shown activity in: U2OS, MCF7, and MDA-MB-231. Based on previous work, U2OS is a cell line with robust circadian rhythms,^31^ the low-grade breast cancer cell line MCF7 has low amplitude oscillations, and the high tumor grade breast cancer cell line MDA-MB-231 is arrhythmic.^32^ We were particularly interested in determining whether the extent of oscillation robustness and nobiletin-influenced circadian changes could be correlated with anti-oncogenic effects. To assess this, we dosed the cells with two concentrations of nobiletin and evaluated how circadian rhythms are affected using real-time luminometry. We assessed the oncogenic features of cell migration and anchorage-independent clonal expansion using wound healing and colony formation assays, respectively. Upon nobiletin treatment, we found subtle circadian alterations in U2OS and MCF7 cells, but strikingly observed enhanced rhythmicity in otherwise arrhythmic MDA-MB-231 cells.^32^ Under the same conditions, oncogenic features in U2OS cells were disparately affected, with motility reduced, but increased colony areas observed. In MCF7 cells, cell migration was not altered, but at the highest concentration, colony areas decreased. The most significant changes were observed in nobiletin treated MDA-MB-231 cells, which possessed reductions in both cell migration and colony area.

## Materials and Methods

### Cell Culture

U2OS cells were obtained from Prof. Patricia Wadsworth (Biology, UMass Amherst), MCF7 cells were obtained from Prof. D. Joseph Jerry (Veterinary and Animal Sciences, UMass Amherst), and MDA-MB-231 cells were obtained from Prof. Shelly Peyton (Chemical Engineering, UMass Amherst). U2OS cells were maintained in DMEM (Gibco), with 10% Fetal Bovine Serum (FBS; Corning), 1% L-Glutamine (Gibco), 1% penicillin-streptomycin (Gibco), 1% Non-Essential Amino Acids (HyClone), and 1% Sodium Pyruvate (Gibco). MCF7 and MDA-MB-231 cells were maintained in DMEM (Gibco), supplemented with 10% FBS (Corning), 1% penicillin-streptomycin (Gibco), and 1% L-glutamine (Gibco). All cells were incubated at 37 °C under 5% CO_2_ atmosphere, unless otherwise noted.

### Lentiviral Transductions

The generation of *Bmal1:luc* and *Per2:luc* plasmids^33^ and their subsequent stable transfection into U2OS,^34^ MCF7, and MDA-MB-231^32^ cells have been described previously.

### Nobiletin Treatments

Nobiletin (Acros Organics) was prepared in 100% dimethyl-sulfoxide (DMSO; Sigma-Aldrich) at a concentration of 191 mM and stored at −20 °C in single use aliquots. When dosing cells, the nobiletin stock was serially diluted in DMSO to achieve the desired concentration needed in order to maintain a final, constant DMSO concentration of 0.2% (including vehicle controls) in media. Media, prepared according to experiment type, containing DMSO/nobiletin was added to synchronized cells (described below).

### Cell Synchronization

Cells were seeded in 35 mm culture dishes in 2 mL at a density of 2 × 10^5^ cells/mL and incubated for 2-4 days (depending on time taken to reach confluence). Synchronization methods differed by cell line based on previous optimizations. U2OS growth media was then aspirated and 100 nM Dexamethasone-containing culture media added for 2 h to synchronize the cells. For MCF7, cells were washed with PBS and synchronized by subjecting to starvation conditions (DMEM with 1% L-glut) for 18 h followed by either treatment with 100 nM Dexamethasone-containing DMEM with 1% L-glut media (for luminometry) or 2 h serum shock in 1:1 FBS and DMEM with 1% L-glut media (for wound healing and colony formation assays). For MDA-MB-231, cells were washed with PBS and synchronized via 18 h starvation in DMEM with 1% L-glut media followed by 2 h serum shock in 1:1 FBS and DMEM with 1% L-glut media. Cells were then treated according to experiment-specific procedures, described below.

### Bioluminescence Recording and Analysis

Following synchronization as described above, media was replaced with bioluminescence recording media. For U2OS cells, bioluminescence recording media was prepared by dissolving powdered DMEM (Sigma-Aldrich) in Millipore-purified water (18.2 MΩ.cm resistance) to give a final concentration of 0.01125 g/mL. This solution was sterile filtered using a 0.2 µm filter (ThermoFisher) and used to prepare recording media containing the following additives (final concentrations are indicated here): 4mM sodium bicarbonate (Gibco), 5% FBS (Corning), 1% HEPES (HyClone), 0.25% penicillin-streptomycin (Gibco), and 150 µg/mL d-luciferin (ThermoFisher). For MCF7 and MDA-MB-231 cells, recording media was prepared by dissolving powdered DMEM (Sigma-Aldrich) in Millipore water to give a final concentration of 0.0135 g/mL. This solution was sterile filtered using a 0.2 µm filter (ThermoFisher) and used to prepare recording media containing the following additives (final concentrations are indicated here): 1% Sodium pyruvate (ThermoScientific), 5% FBS, 1% HEPES, 1% penicillin-streptomycin, and 150 µg/mL d-luciferin (ThermoFisher). Dishes were sealed with 40 mm sterile cover glass using silicon vacuum grease and subjected to monitoring using a LumiCycle 32 System (Actimetrics) at 36.5 °C for 5-7 days.

Each bioluminescence recording was pre-processed to remove an initial 12 h transient and spikes. To assess the rhythmicity of each recording, a linear trend was removed, and then an FFT-based rhythmicity test^35^ was applied. Rhythmic time-series were de-trended by removing the 24 h moving average (which excludes an additional 12 hours of data) and smoothed using the 3 h running average method. Circadian amplitude, period, and damping rate of each time-series were estimated with two approaches. The first was to fit the first three cycles (t=24 h to t=96 h) to a damped cosine curve with a nonlinear least squares method, using R (www.r-project.org) code adapted from Hirota et al.^36^ The second approach was to identify the time and magnitude of each phase marker (peak, trough, mean-crossing on the rise, mean-crossing on the fall) of the first three cycles (custom Matlab script). Each phase marker was used to estimate a period (e.g. average peak-to-peak measurements led to one estimate and trough-to-trough to a second), providing 4 additional period estimates. The amplitude was estimated by adding the magnitude of the first trough to the magnitude of the first peak. To capture damping, the ratio of the amplitude of the second cycle to that of the first cycle was subtracted from unity.

### Wound Healing Assay

U2OS, MCF7 and MDA-MB231 cells were seeded in 0.5 mL at a density of 2×10^5^ cells/mL (U2OS) and ~4×10^5^ cells/mL (MCF7, MDA-MB-231), in 24-well plates and incubated until 100% confluence was reached (2-4 days). Cells were synchronized as described above. Wounds were generated using a 1 mL micropipette tip. Culture media was then removed, cells were washed with PBS (Gibco), and 500 µL of new culture media containing indicated treatments was added into each well. Images were taken immediately following for the first time point (T=0) and then every 2 h for 24 h via Biotek Cytation 3 cell imaging multi-mode plate reader. Wound closure (%) was determined by normalizing wound area quantified at each time point via macros adapted from MRI wound healing tool (http://dev.mri.cnrs.fr/projects/imagej-macros/wiki/Wound_Healing_Tool) to T=0 via ImageJ.

### Colony Formation Assay

Cells were suspended in agarose and incubated until colonies formed, similarly to a previously described procedure.^33^ Briefly, 3% 2-Hydroxyethyl Agarose (Sigma) was prepared and stored in a water bath at 45 °C. Cell culture media at 37 °C was used to dilute agarose to 0.6%, and 500 µL of the resulting solution was added to each well of a 24-well plate (Nuclon), which was incubated at 4 °C until the agarose solidified to produce the first layer. The agarose solution was also added to the suspended cells (U2OS: 1 × 10^4^ cells/mL; MCF7 and MDA-MB-231 3.5 × 10^3^ cells/mL) in warm culture media to 0.3%, followed by addition of nobiletin. 500 µL of the drug-cell-agarose solution was dispensed per well and incubated at 4 °C for 30 min until the second agarose layer solidified. Then plates were transferred and incubated at 37 °C under 5% CO_2_. A feeder layer of 0.3% agarose in culture media containing nobiletin at designated concentrations (in 500 µL) was added to each well once every 7 days for 4 weeks. When imaging colonies, 20% methanol (Fisher Scientific) was added to wells, and plates were incubated on a shaker at rt for 1 h. Eight images were taken per well using Biotek Citation 3 multi-mode cell imaging reader. Each condition was performed in three biological replicates. Images were stitched and colony numbers/sizes were analyzed using ImageJ software.

## Results

### Nobiletin affects circadian oscillations of *Per2:luc* and *Bmal1:luc* in U2OS, MCF7, and MDA-MB-231 cells

To assess the circadian effects of nobiletin in a detailed manner, we used luciferase-reporter cells for *Bmal1* and *Per2* promoter activity previously generated in our lab.^33^ Here, we employed U2OS, MCF7 and MDA-MB-231 cell lines separately transfected with *Bmal1:luc* and *Per2:luc.*^32,34^ Prior to initiating luminometry experiments, we evaluated potential viability effects of treatments to be used. The highest and lowest concentrations used in this study were determined based on the concentration required for the 50% induction of activity in MCF7 cells (44 µM).^37^ Therefore, concentrations of 5 µM and 50 µM of nobiletin were used. While slightly diminished viability (~80%) was observed for U2OS cells treated at the highest concentration, none of the other cells or treatments resulted in significant changes (Supplementary Fig. S1). The same conditions were used to track circadian oscillations via reporter bioluminescence using real-time luminometry for 6 days (Fig. 1, S2, S3, S4, S5). A total of N=12 replicates per condition per cell line/reporter was obtained, across three independent experiments, with N=4 each. We observed that consistent, rhythmic, anti-phase oscillations are observed for both *Bmal1:luc and Per2:luc* signals in U2OS and MCF7 cells across all treatments. However, in MDA-MB-231 cells, previously reported to lack detectable rhythms,^32^ *Bmal1* and *Per2* reporters showed oscillations consistently only with 50 µM nobiletin treatment. We also confirmed that the treatment of cells with DMSO vehicle did not affect circadian oscillations, in comparison with non-treated cells (Supplementary Fig. S6, S7).

**Figure 1.**
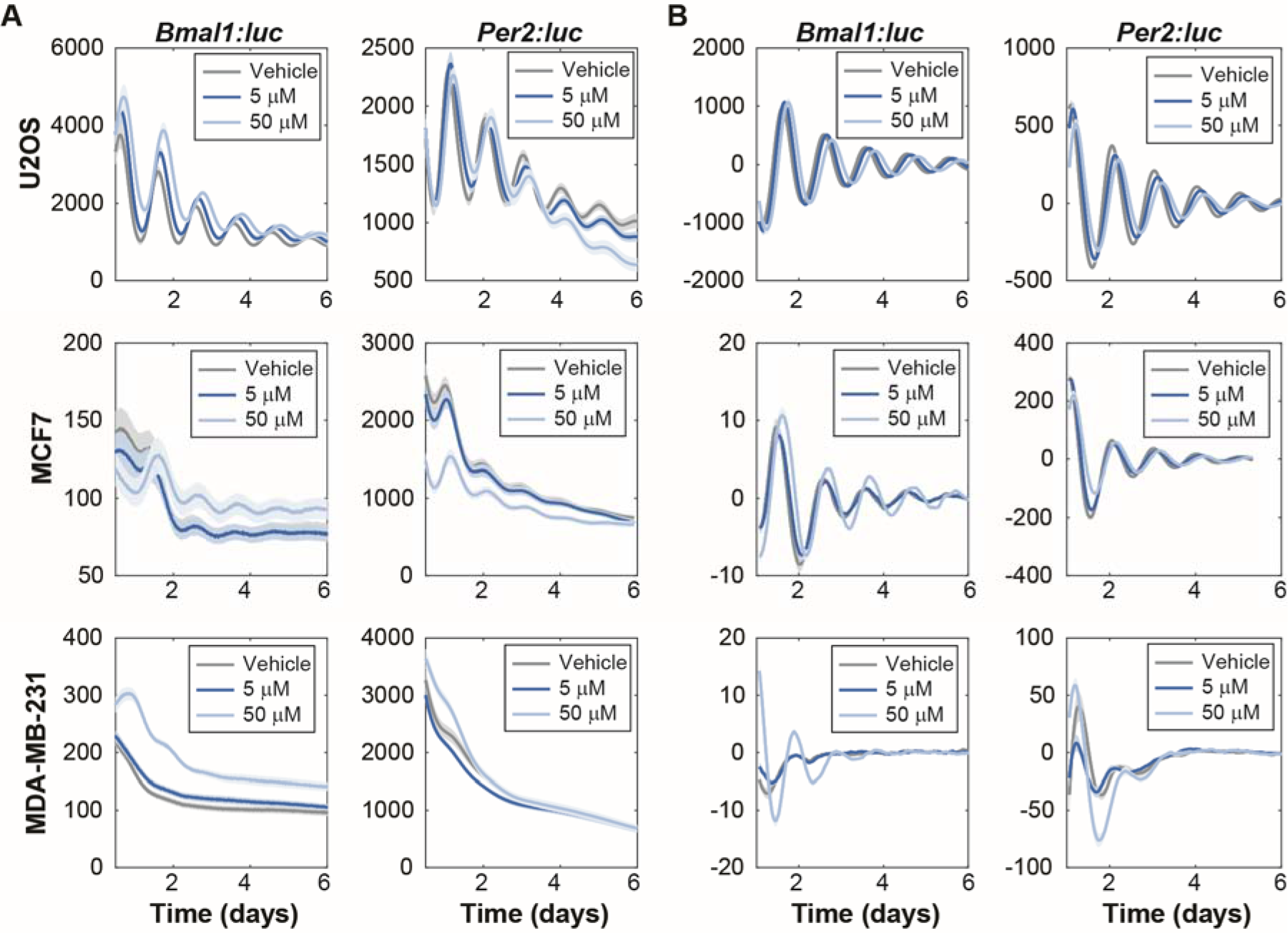
Nobiletin alters circadian dynamics in bone and breast cancer cell lines. Shown are **(A)** raw and **(B)** de-trended mean traces of (left) *Bmal1:luc* and (right) *Per2:luc* in (top) U2OS, (middle) MCF7, and (bottom) MDA-MB-231 cells, as obtained using luminometry. Each cell type was exposed to DMSO (vehicle), and 5 μM and 50 μM nobiletin conditions. The semi-transparent envelopes around the lines indicate the SEM. In several instances, the SEM is too small to extend beyond the line. **(B)** Each trace was de-trended by subtracting the mean of a 24-h sliding window and smoothed using the mean of a 3-h sliding window.

We further evaluated the characteristics of data from U2OS and MCF7 cells (MDA-MB-231 cells were omitted due to lack of rhythmic patterns and are discussed further below). Our analyses show that nobiletin results in modestly increased circadian periods in both U2OS-*Bmal1:luc* and U2OS-*Per2:luc* cells (Fig. 2), with higher concentrations yielding greater effects (randomization test for difference in means, Bonferroni-corrected p < 0.05). The changes to damping rates and amplitudes were not statistically significant, with the exception that 5 µM nobiletin treatment led to more damped *Per2:luc* oscillations. A repeated analysis with peak- and trough-finding methods for estimating the period, amplitude, and damping rate led to the same trends (Supplementary Fig. S8, S9, S10, S11). For MCF7 cells, the trends in period, amplitude, or damping rate estimates across treatments varied, depending on the method used to estimate the properties (Supplementary Fig. S8, S9, S10, S11).

**Figure 2.**
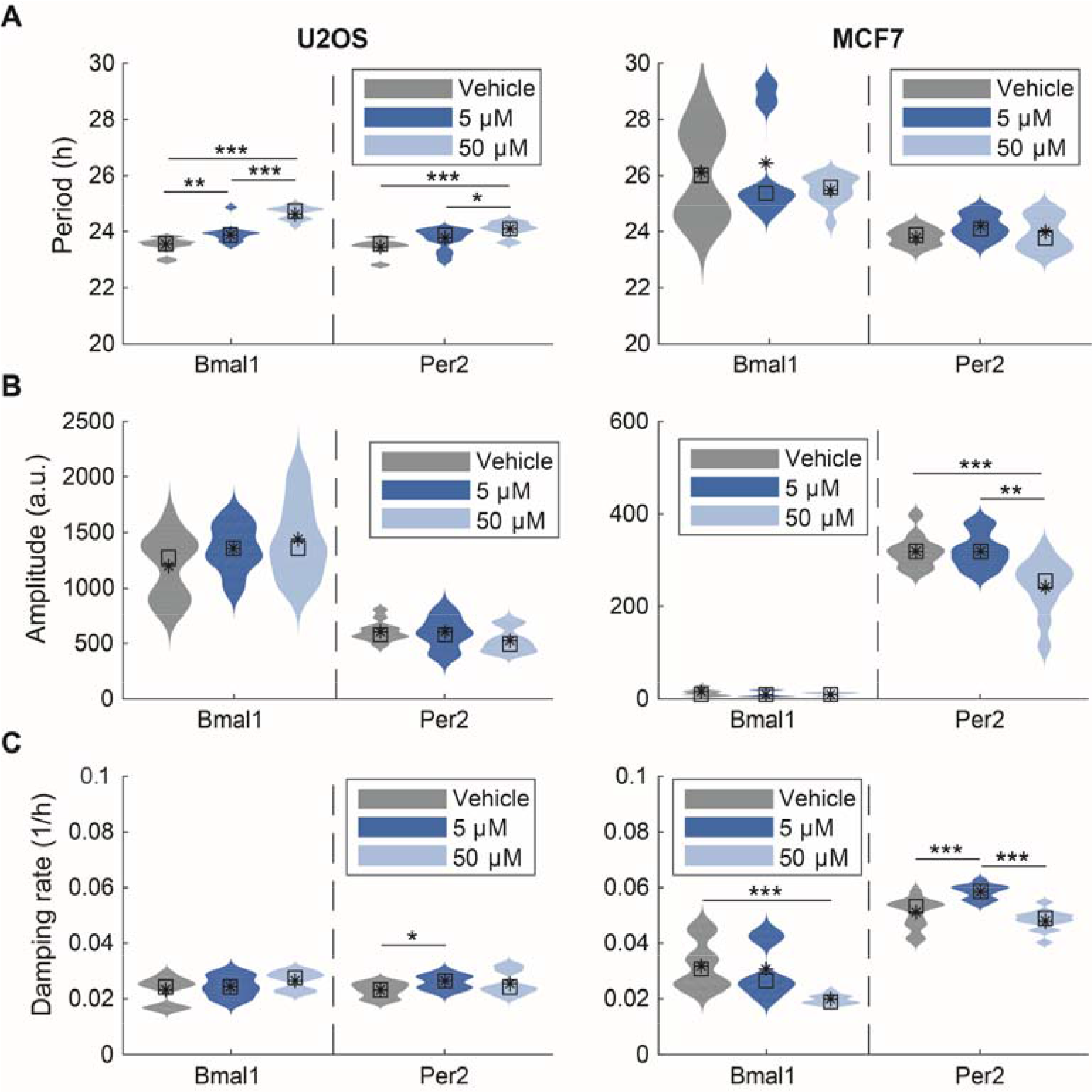
Effects of nobiletin on circadian periods, amplitudes, and damping rates in (left) U2OS and (right) MCF7 cell lines. Shown are the distributions of **(A)** period, **(B)** damping rate, and **(C)** amplitude parameters estimated from fitting a damped cosine curve to each de-trended bioluminescence recording. Within each subfigure, color indicates experimental condition, with results from *Bmal1:luc* shown on left and *Per2:luc* on the right (N=12 per reporter per condition). The mean and median values are indicated with a black asterisk and square, respectively. Statistical significance was evaluated via a randomization test for difference in means with a Bonferroni correction (*p < 0.05, **p < 0.01, ***p < 0.001),

### Nobiletin affects rhythmicity of MDA-MB-231 cell lines

Previous work by our group has shown that even using real-time luminometry with *Bmal1* and *Per2* reporters, circadian oscillations remain undetectable (and arrhythmic) in MDA-MB-231 cells.^32^ We find that this largely remains the case, however, at the highest concentration of nobiletin (50 µM), rhythmicity was found to be enhanced in both *Bmal1:luc* and *Per2:luc* cells. This was determined by applying an FFT-base test for rhythmicity to recordings after removing an exponential (Fig. 3) or quadratic (Supplementary Fig. S12) trend. The results varied depending on the length of time-series considered, but recordings from cells treated with 50 µM nobiletin consistently scored rhythmic at a higher rate than all others. Visual inspection of the de-trended time-series confirms that nobiletin enhances rhythmicity (Supplementary Fig. S13, S14, S15, S16). To assess whether nobiletin resulted in enhanced transcription of *Bmal1* or *Per2*, we conducted an RT-PCR experiment with MDA-MB-231, U2OS, and MCF7 cells (Supplementary Fig. S17). We found that no significant changes occurred following nobiletin treatment across all of the cell lines.

**Figure 3.**
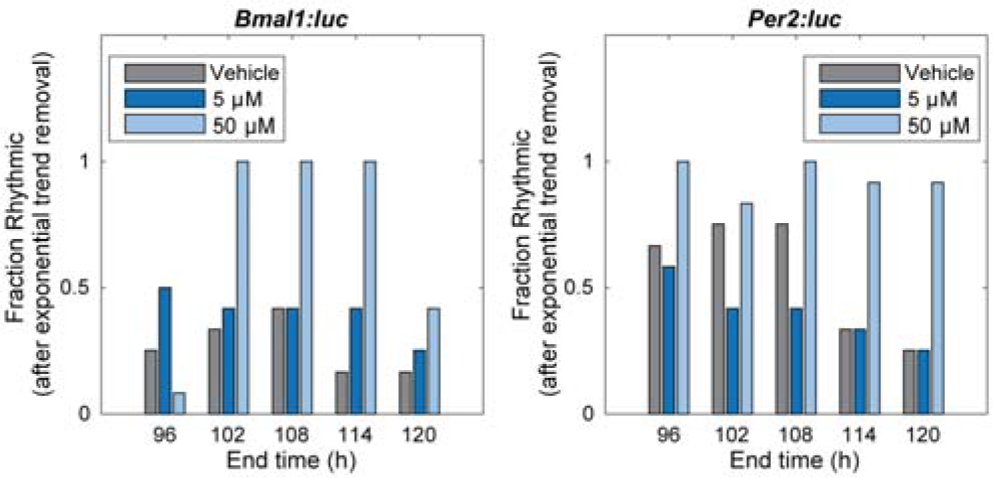
Nobiletin alters the rhythmicity of MDA-MB-231 cells. Shown are the fractions of recordings classified as rhythmic (p < 0.05 on FFT-based test), after removing an exponential trend, using time-series with increasing end times. The fraction scored rhythmic depends on how much of the time-series is included in the analysis, but cells treated with 50 μM nobiletin consistently scored as rhythmic more frequently than those treated with either DMSO or 5 μM. The only exception is *Per2:luc* recordings with the first 108 hours analyzed.

### Effects of nobiletin on oncogenic characteristics vary by cell type

Previous studies showed that nobiletin inhibits the proliferation of U2OS, MCF7 and MDA-MB-231 cells^23,26, 37^ and reduces their migration and invasion.^22,23,26,30^ As nobiletin has been posited to be a compound with therapeutic potential, we were interested to determine whether its effects were similar or differed by cell line. First, we assessed changes to cellular migration following nobiletin treatment via wound-healing/scratch assay (Fig. 4). The same concentrations were used as in our assessments of circadian rhythms. In U2OS and MDA-MB-231 cells, migration substantially decreased in samples treated with 50 μM nobiletin. However, in MCF7 cells, nobiletin had no significant effect on cell migration.

**Figure 4.**
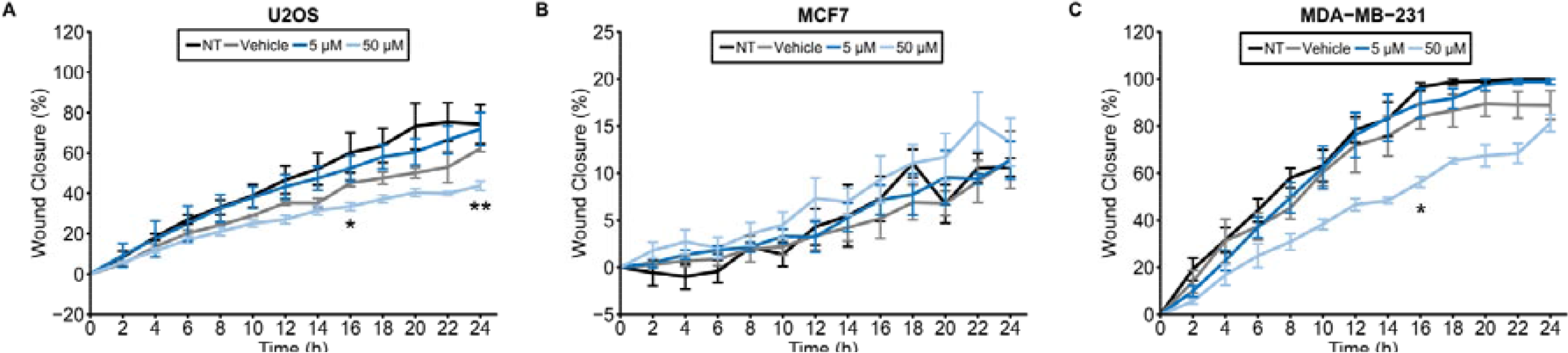
Effects of nobiletin on cellular migration of bone and breast cancer cells. A wound healing/scratch assay was performed with (**A**) U2OS, (**B**) MCF7, and (**C**) MDA-MB-231 cells. Nobiletin significantly reduced cell migration in U2OS and MDA-MB-231 cell lines at 50 µM concentration. No substantial changes were observed in MCF7 cells. Error bars represent standard error of the mean (SEM). Statistical significance was evaluated via Student’s T.test, with Bonferroni correction (*p < 0.05, **p < 0.01). NT = non-treated; Vehicle = DMSO-only control (0.2%).

We used a colony formation assay to determine whether nobiletin could affect anchorage-independent colony formation in a three-dimensional environment. In this experiment, we evaluated both colony area and size. In U2OS cells, colony areas significantly increased (Fig. 5A, S18), however the numbers of colonies present did not change compared to the vehicle-treated samples (Fig. 5D). In MCF7 cells, nobiletin significantly reduced the area of colonies (Fig. 5B, S18), but did not alter their number (Fig. 5E). However, the most substantial change in colony size was observed in MDA-MB-231 cells (Fig. 5C, S18), which also showed increased colony numbers (Fig. 5F). Decreased colony sizes may be accompanied by increased numbers because it is possible that nobiletin prevents the aggregation of smaller colonies into larger ones. This can result in higher numbers of smaller colonies but fewer larger ones.^34^

**Figure 5.**
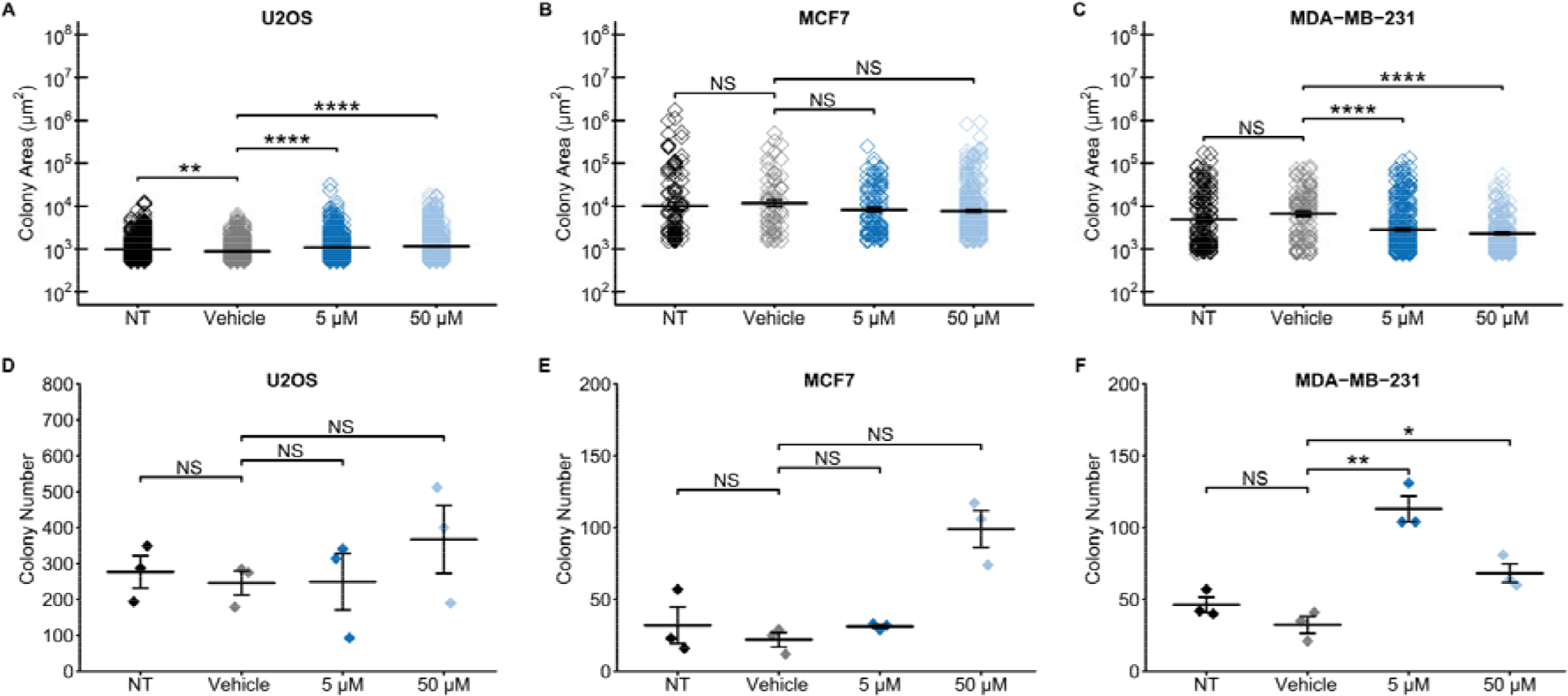
Nobiletin alters colony area and number of bone and breast cancer cells. The colony formation assay was performed in (**A**) U2OS, (**B**) MCF7, and (**C**) MDA-MB-231 cells. Nobiletin increased colony area in U2OS cells in a statistically significant manner, decreased the colony area at 50 µM in MCF7 cells, and significantly reduced colony area in a dose dependent manner in MDA-MB-231 cells. However, the colony numbers were not affected in (**D**) U2OS **(E)** MCF7 cells, and significantly altered at both treatments in (**F**) MDA-MB-231 cells. Error bars represent standard error of the mean (SEM). Statistical significance was evaluated via Student’s T test with a Bonferroni correction (*p < 0.05, **p < 0.01, ***p < 0.001, ****p < 0.0001). NT = non-treated; Vehicle = DMSO-only control (0.2%), NS = non-significant.

## Discussion

Nobiletin is a small molecule with a range of therapeutic effects whose influence on circadian rhythms has recently been uncovered. While numerous studies have evaluated its efficacy in various disease models, to our knowledge only one assessed therapeutic and circadian effects concomitantly, addressing metabolic syndrome.^2^ Because nobiletin has been shown to result in beneficial outcomes in multiple cancer types, we considered it important to address whether a correlation may exist between its anti-oncogenic and circadian changes. Circadian rhythm disruption has been found to be associated with cancers and their aggressiveness. As examples, peripheral blood mononuclear cells (PBMCs) from healthy individuals had circadian oscillations while chronic myeloid leukemia did not,^38^ low tumor grade MCF7 cells had circadian rhythms while high tumor grade MDA-MB-231 cells were arrhythmic,^32^ and in models of disease progression circadian discrepancies were observed in both breast,^33^ and colon carcinoma cells.^39^ In this study, we assessed how nobiletin affects circadian and cancerous characteristics in three different cell lines. We hypothesized that nobiletin’s effects on oncogenic features, including motility and anchorage-independent colony formation, would be accompanied by changes in circadian rhythms, and that their extents may be linked. We employed U2OS, MCF7, and MDA-MB-231 cells; while the first two possess strong and weaker oscillations, respectively, MDA-MB-231 cells had been previously found to be arrhythmic.^32^ Following treatment with nobiletin, U2OS and MCF7 showed subtle circadian alterations and slight changes to cellular characteristics. In contrast, the MDA-MB-231 responses included a clear enhancement of circadian oscillations and significant reduction in cell migration and colony area.

Previous studies using clock knockout and normal MEF cells showed amplitude enhancements upon nobiletin treatment.^2,3^ These effects are hypothesized to be the result of nobiletin’s activation of the BMAL1 transcription factor ROR. In our work, U2OS-*Bmal1:luc*, and MDA-MB-231 *Bmal1:luc* and *Per2:luc* cell lines showed amplitude enhancements, with rescue of rhythmicity in MDA-MB-231 cells. However, no amplitude changes were observed in U2OS-*Per2:luc* cells and nobiletin subtly reduced circadian amplitudes of both MCF7 *Bmal1:luc* and *Per2:luc* cell lines. Our raw data supports these findings, with the exception of MCF7 *Bmal1:luc*, where bioluminescence was enhanced. Nobiletin’s enhancement of circadian amplitude may vary depending on initial characteristics. For example, significant amplitude enhancement was observed previously in MEFs where CLOCK was knocked-out,^2^ while standard MEFs showed more minor effects.^3^ It is possible that the outcomes of nobiletin treatment are dependent on circadian (dys)function. We observed that rhythmic U2OS and MCF7 cells showed subtle changes, while arrhythmic MDA-MB-231 cells showed clear alterations with nobiletin. Those cell lines where the clock was clearly functioning benefitted less than the one where it was not. In terms of period, prior work showed that nobiletin increases the period of PER2::LUC MEFs,^2,3^ and mouse liver slices.^3^ We observed similar but subtle effects in both *Bmal1* and *Per2* U2OS reporter cells. However, MCF7 cells had no significant changes in period. In MDA-MB-231 cells, periods could not be determined for arrhythmic oscillations, but for rhythmic ones, periods were shown to be within the circadian range. Our data showed different circadian effects between the two reporters associated with the same cell line to same nobiletin treatments (e.g. the periods of U2OS-*Bmal1:luc* versus -*Per2:luc* and the periods and amplitudes of MCF7-*Bmal1:luc* versus *-Per2:luc* cell lines). Such disparate responses from reporters in the same cell type have been seen previously in U2OS and NIH 3T3 cell lines.^40^ This could be a result of having each reporter in a separate cell line. Therefore, in the future, tracking both *Bmal1* and *Per2* rhythms concomitantly using a dual-luciferase reporter system will be helpful.

Nobiletin also affected oncogenic features of the cancer cells to varying extents. Sheet migration is a process where cancer cells migrate in two dimensions during metastasis, and can be assessed via wound healing assay.^41^ Furthermore, normal cells rely on contacts between cells and extra cellular matrix for cell growth and division, while more aggressive ones can grow without it. The ability of this anchorage-independent cell proliferation can be measured using a colony formation assay.^42^ The colony counts and areas obtained from this assay are quantitative values that can determine tumorigenicity of a cell line.^43^ In this study, we measured the ability of cancer cells to migrate and proliferate in an anchorage-independent manner in three dimensions^42^ using both of these methods. We saw significantly reduced cell migration at the highest concentration of nobiletin treatment in U2OS and MDA-MB-231 cells, but not in MCF7 cells. Some of the pathways associated with anti-oncogenic activities in the three cell lines tested are connected to the circadian clock. A previous study showed that nobiletin reduced cell migration (to a similar extent) via downregulation of MMPs and nuclear NF-κB protein levels in U2OS cells.^26^ MMPs are repressed by the circadian protein-PER,^44^ while NF-κB has been shown to be downregulated by ROR.^45^ Hence, ROR activation and enhanced clock activity may affect migration. Our data shows that nobiletin also reduced MDA-MB-231 migration at the highest concentration tested; a previous study (using a much higher concentration, 200 µM) showed that nobiletin inhibits motility by affecting CD36, pSTAT3, and NF-κB proteins.^30^ STAT3 has shown circadian rhythms in rat SCN,^46,47^ and it is possible that nobiletin affects its oscillations as well. In our study, nobiletin did not alter MCF7 cell migration at the highest concentration tested (50 µM). However, previous studies showed that it reduced cell migration at concentrations of 50 µM,^22^ 100 µM and 200 µM.^23^ Multiple experiments showed that nobiletin downregulates CD36, STAT3, pSTAT3, MMPs, and NF-κB proteins, among others in MCF7 cells.^22,23,30^

Colony areas were concomitantly increased and decreased with dose in U2OS and MDA-MB-231 cells, respectively, while MCF7 was affected only at the highest concentration used. Previous studies showed that nobiletin-treated breast cancer cells had reduced cell proliferation in largely two-dimensional assays.^24,37^ In one, nobiletin caused G1 cell cycle block in MCF7 cells at 75 μM and 100 μM concentrations,^37^ while in another it arrested MCF7 cells in the G0/G1 phases of the cell cycle at 50 μM and decreased ERK activation and suppressed cyclin D1 expression.^48^ Cyclin D1 is a key regulator of G_0_/G_1_ cell-cycle checkpoint, and is under circadian regulation by a PER1-protein complex.^49^ Hence, circadian enhancement via nobiletin may be involved. Another study using MCF7 and MDA-MB-231 cells also showed that nobiletin reduced cell proliferation in the concentration range of 50 µM-300 µM, and affected the SRC/FAK/STAT3 pathway.^23^ To the best of our knowledge, nobiletin has been evaluated only in terms of three-dimensional growth in MCF7 cells using a sphere formation assay, where significant reductions in size were observed at 200 µM.^30^ We also found decreased colony areas at the highest concentration tested. Colony formation assays in other cancer cell lines such as ACHN and Caki2, showed that nobiletin dose-dependently reduced colony numbers at concentrations up to 120 µM,^18^ and colony numbers of H460 non-small cell lung cancer cells at 50 µM.^50^

Interestingly, in our study, we found that nobiletin yields different circadian and anti-cancer effects depending on cell line. Another significant difference among these cells is the levels of nobiletin metabolizing protein, Cytochrome P450 (CYP), present. CYP metabolizes naturally occurring flavonoids; the metabolized products often have higher activities compared to parent compounds.^37,50^ Nobiletin has been shown to be metabolized via CYP1.^24,37^ While CYP1 protein levels in U2OS cells have not yet been determined, MDA-MB-231s have higher CYP1 expression compared to MCF7.^51^ While the IC_50_ of nobiletin has not been determined in MDA-MB-231 cells, in another triple negative cell line where CYP1 was shown to be constitutively expressed, MDA-MB-468,^24^ the IC_50_ of nobiletin is 0.1 ±0.04 μM.^24^ In contrast, in MCF7 cells it is 44 μM.^37^ In the future, it would be useful to determine whether nobiletin’s metabolites are responsible for its activity, and to directly compare its effects in these cell lines in parallel.

We conclude that alteration to cancer cell circadian rhythms and oncogenic features via nobiletin treatment occur concomitantly. We have observed that cells with greater circadian changes also have more substantial effects on cellular characteristics, and vice-versa. In the future, it is highly merited to study whether there is a direct relationship between circadian alterations (rescue/enhancement) and affecting oncogenic features in cancer cells.

## Acknowledgements

We are grateful to Tanya Leise (Amherst College Mathematics) for helpful discussions. HHL was funded by a UMass Amherst Chemistry-Biology Interface Training Program Fellowship; KLR was funded by a UMass Honors Research Grant; SRT was funded by the Office of the Provost at Colby College; MEH was funded by the NIH (R15 GM126545). We are also grateful to the laboratory of Yubing Sun (UMass Amherst Chemical Engineering) for plate reader access.

## Author Contributions

MEF and MEH conceived of the studies; SSLD, KLR, HHL, and CL performed experiments; SSLD and SRT performed data analyses; SSLD, SRT, and MEF generated all figures; SSLD, KLR, CL, SRT, and MEF wrote the manuscript. All authors reviewed and revised the manuscript.

## Data Availability

The datasets generated during and/or analyzed during the current study are available from the corresponding author on reasonable request.

