## Supporting Information for "Nobiletin Affects Circadian Rhythms and Oncogenic Characteristics in a Cell-Dependent Manner"

### Methods

### Cell Viability Assay

U2OS, MCF7, and MDA-MB-231 cells were seeded in a 24 well plate in 0.5 mL at a density of $2\times10^5$ cells/mL, and grown to confluence. Depending on sample, cells were not treated, or treated with DMSO vehicle (to 0.2%), 5 µM nobiletin or 50 µM nobiletin (both with 0.2% DMSO). Cells were incubated for 24 h at 37 °C in 5% $CO_2$. Then the culture media was replaced with 10% Alamar blue reagent (Invitrogen) and incubated for 1 h under the same conditions. The Alamar blue reagent was then transferred to a black 96 well plate (Costar), and samples excited at 540 nm, and emission detected at 590 nm using a Synergy H1 microplate reader.

### RNA Isolation, cDNA Conversion and Real time PCR

U2OS, MCF7, and MDA-MB-231 cells were seeded in a 24 well plate in 0.5 mL at a density of $2\times10^5$ cells/mL and grown for 2-4 days. Depending on sample, cells were not treated, or treated with DMSO vehicle (to 0.2%), 5 µM nobiletin or 50 µM nobiletin (both with 0.2% DMSO). Cells were then incubated for 24 h. After, the RNA was harvested as described previously.[1] Briefly, RNA was harvested from cells using a PureLink RNA Mini Kit (Ambion) according to manufacturer's instructions. RNA was reverse transcribed to cDNA using 50 µM random hexamers, 10 mM dNTPs, 40 U/µL RNaseOut, and 200 U/µL SuperScript IV Reverse Transcriptase, 100 mM DTT, and 5x SSIV buffer (Thermo Fisher Scientific). RT-qPCR was performed in 96-well plates. Each reaction consisted of 100 ng cDNA, 10 µL iTaq universal SYBR Green Supermix (Biorad), 4 µM each forward and reverse primer (Integrated DNA Technologies), and RNAse-free water to 20 µL. The primers used are *GAPDH* forward (5'- CTT CTT TTG CGT CGC CAG CC-3'), reverse (5'-ATT CCG TTG ACT CCG ACC TTC-3'); *BMAL1* forward (5'- CTA CGC TAG AGG GCT TCC TG-3'), reverse (5'- CTT TTC AGG CGG TCA GCT TC-3'); *PER2* forward (5'- TGT CCC AGG TGG AGA GTG GT-3'), reverse (5'- TGT CAC CGC AGT TCA AAC GAG-3'). The samples were centrifuged and analyzed via CFX Connect real-time system (Biorad) using an initial denaturation at 95°C for 3 min, followed by 40 cycles of 95°C denaturation for 10 s, and 60°C annealing/extension for 30 s. Relative *BMAL1* and *PER2* expression were determined by comparing $C_t$ values of *BMAL1* and *PER2* to *GAPDH* (control) via $2^{\Delta\Delta C_t}$ method.[2] Three biological replicates and three technical replicates per biological replicate were analyzed for each condition.

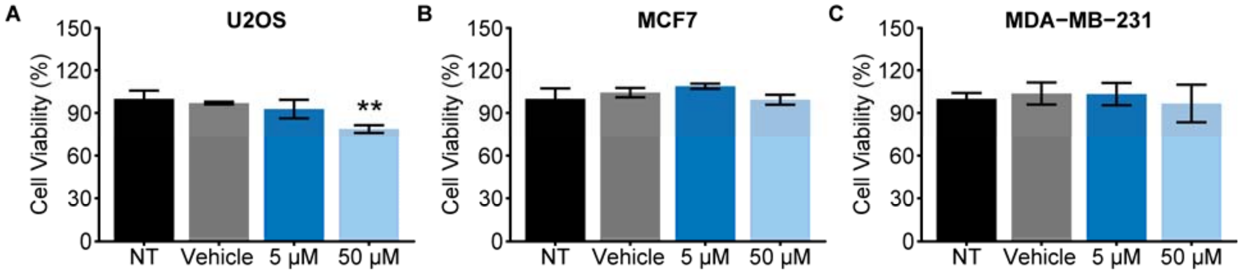


**Fig S1. Cell viability assay. (A)** U2OS, **(B)** MDA-MB-231, and **(C)** MCF7 cells were treated under conditions shown. Viability was determined using Alamar blue assay. No significant differences were observed in nobiletin-treated samples compared to vehicle (0.2 % DMSO) samples, with the exception of the 50 μM U2OS cells. Error bars represent standard deviations across three biological replicates. Statistical significance was evaluated via two tailed Student's T test (** Bonferroni-corrected $p<0.01$). NT = non-treated.

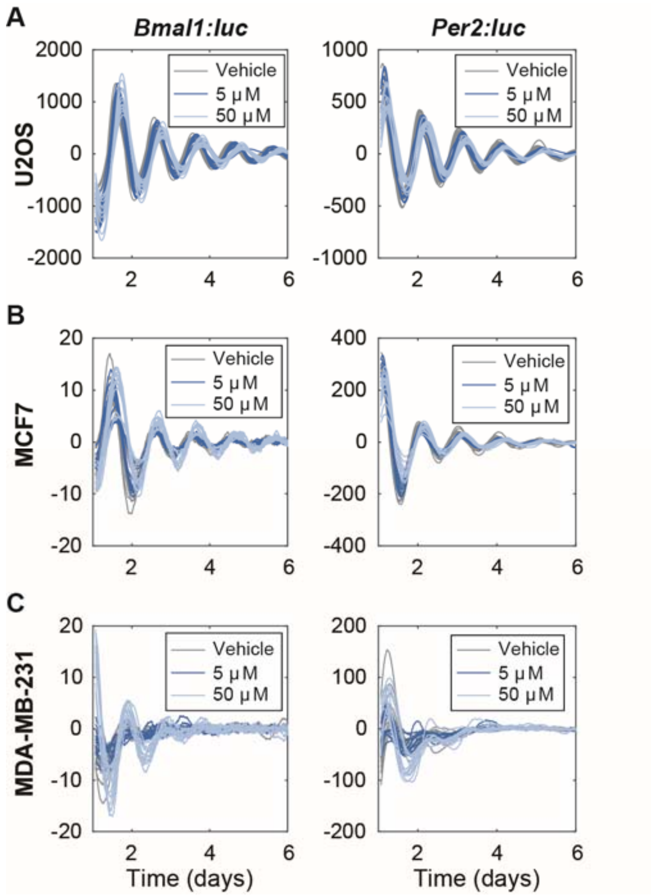


**Fig. S2**. **Effects of nobiletin on circadian dynamics in bone and breast cancer cell lines.** Replicates of the luminometry data shown in **Figure 1** are presented here, for (left) *Bmal1:luc* and (right) *Per2:luc* in (**A**) U2OS, (**B**) MCF7, and (**C**) MDA-MB-231 cells. Each cell line was treated with vehicle (0.2% DMSO), and 5 μM and 50 μM nobiletin conditions. N=12 replicates were obtained for each treatment and cell line. Data were evaluated beginning from t=0.5 (day); each trace has been de-trended by subtracting the mean of a 24h sliding window and smoothed using the mean of a 3-h sliding window, making the de-trended date begin at t=1 (days).

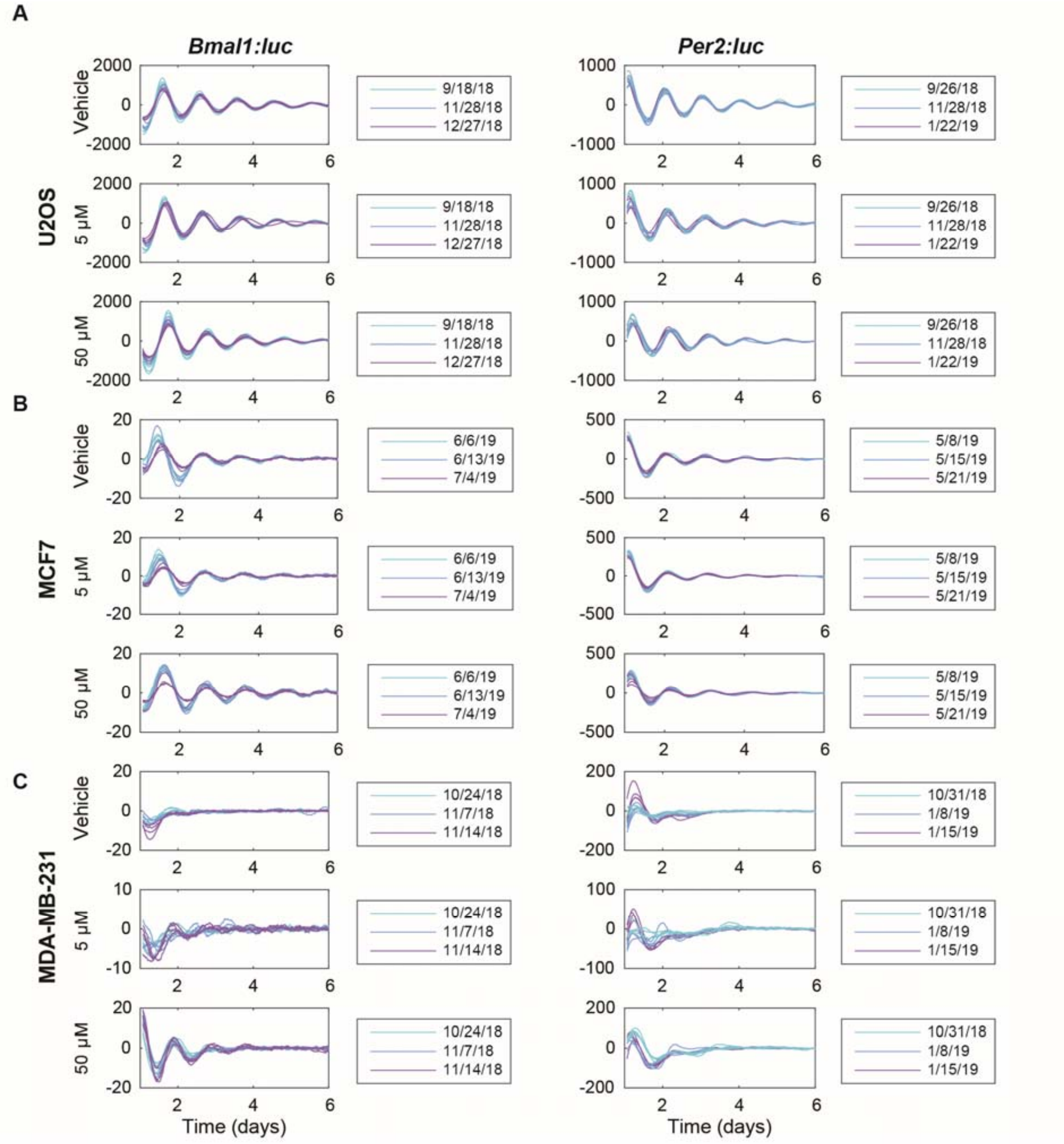


**Fig S3. Effects of nobiletin on circadian dynamics in bone & breast cancer cell lines.** We show the individual replicates depicted in **Fig. S2**, further separated by condition and date. N=4 replicates were obtained for each treatment and cell line, on each of three separate dates (indicated in the legends). Data were evaluated beginning from t=0.5 (day); each trace has been de-trended by subtracting the mean of a 24h sliding window and smoothed using the mean of a 3-h sliding window, making the de-trended date begin at t=1 (days).

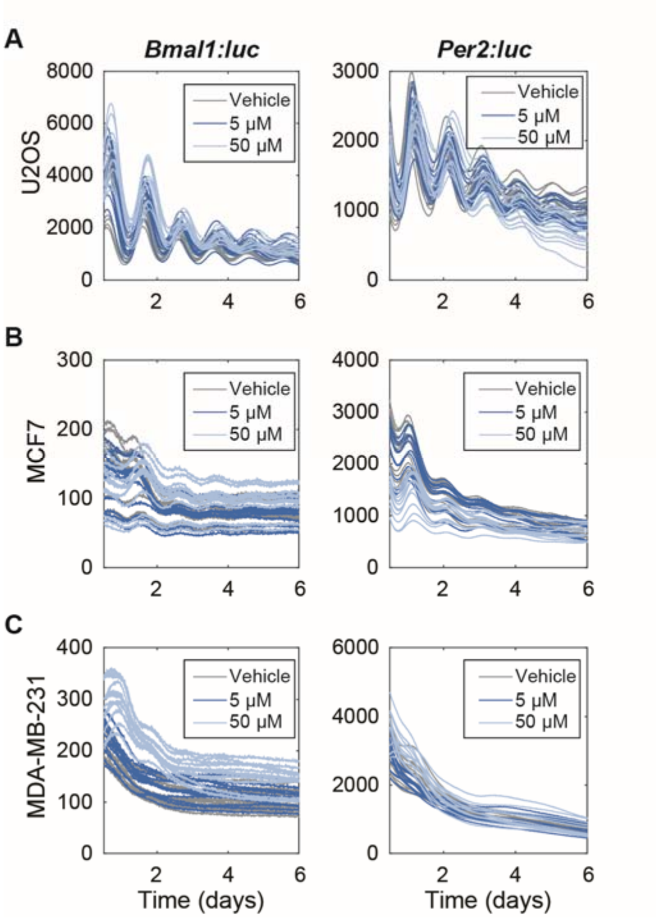


**Fig. S4**. **Effects of nobiletin on circadian dynamics in bone and breast cancer cell lines.** Shown are the raw data for (left) *Bmal1:luc* and (right) *Per2:luc* traces in (**A**) U2OS, (**B**) MCF7, and (**C**) MDA-MB-231 cells under vehicle (0.2% DMSO), and 5 μM and 50 μM nobiletin treatment conditions. N=12 replicates were obtained for each treatment and cell line.

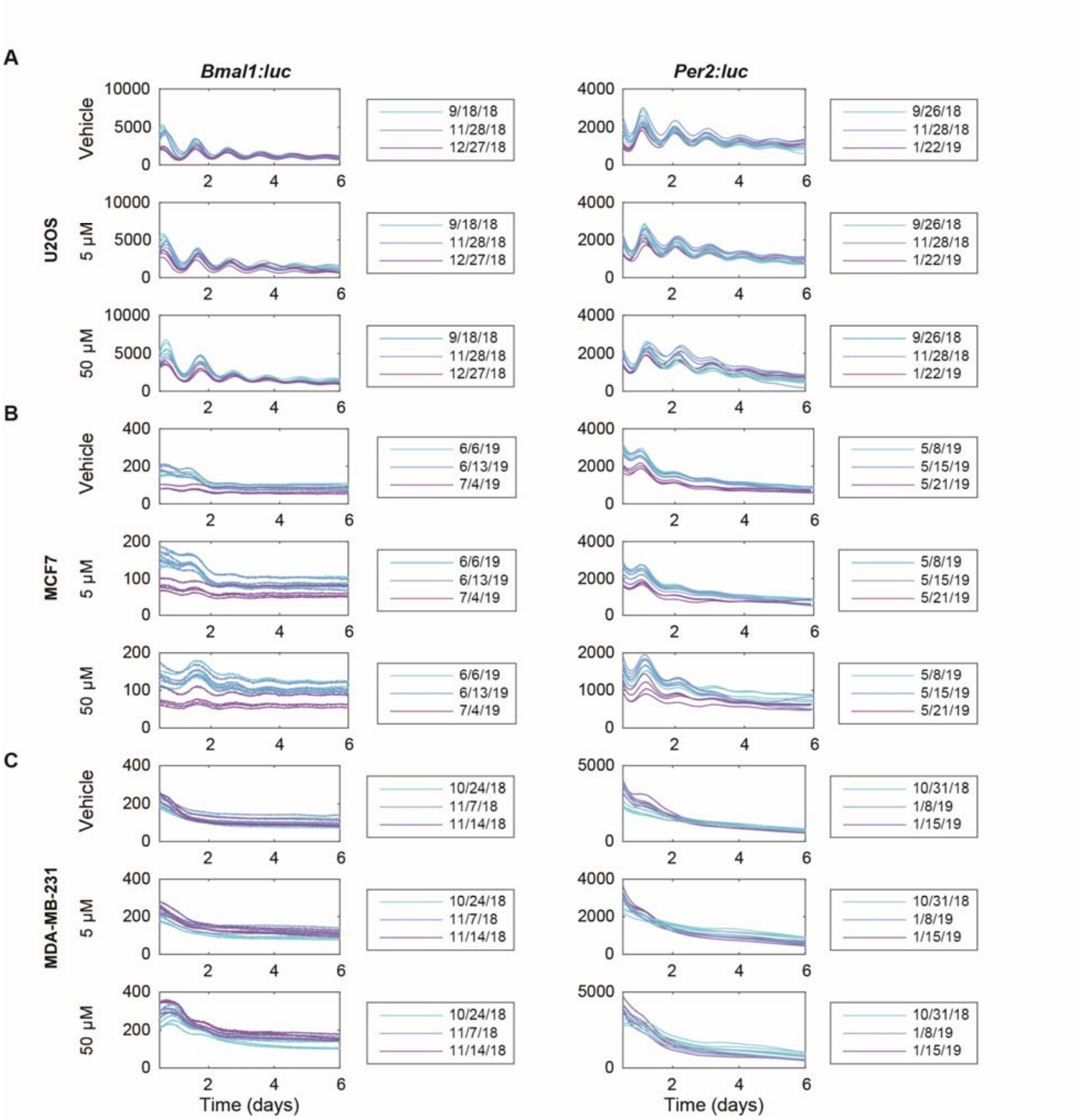


**Fig S5**. **Effects of nobiletin on circadian dynamics in bone and breast cancer cell lines.** We show the individual replicates of raw data depicted in **Fig. S4**, further separated by condition and date. N=4 replicates were obtained for each treatment and cell line, on each of three separate dates (indicated in the legends).

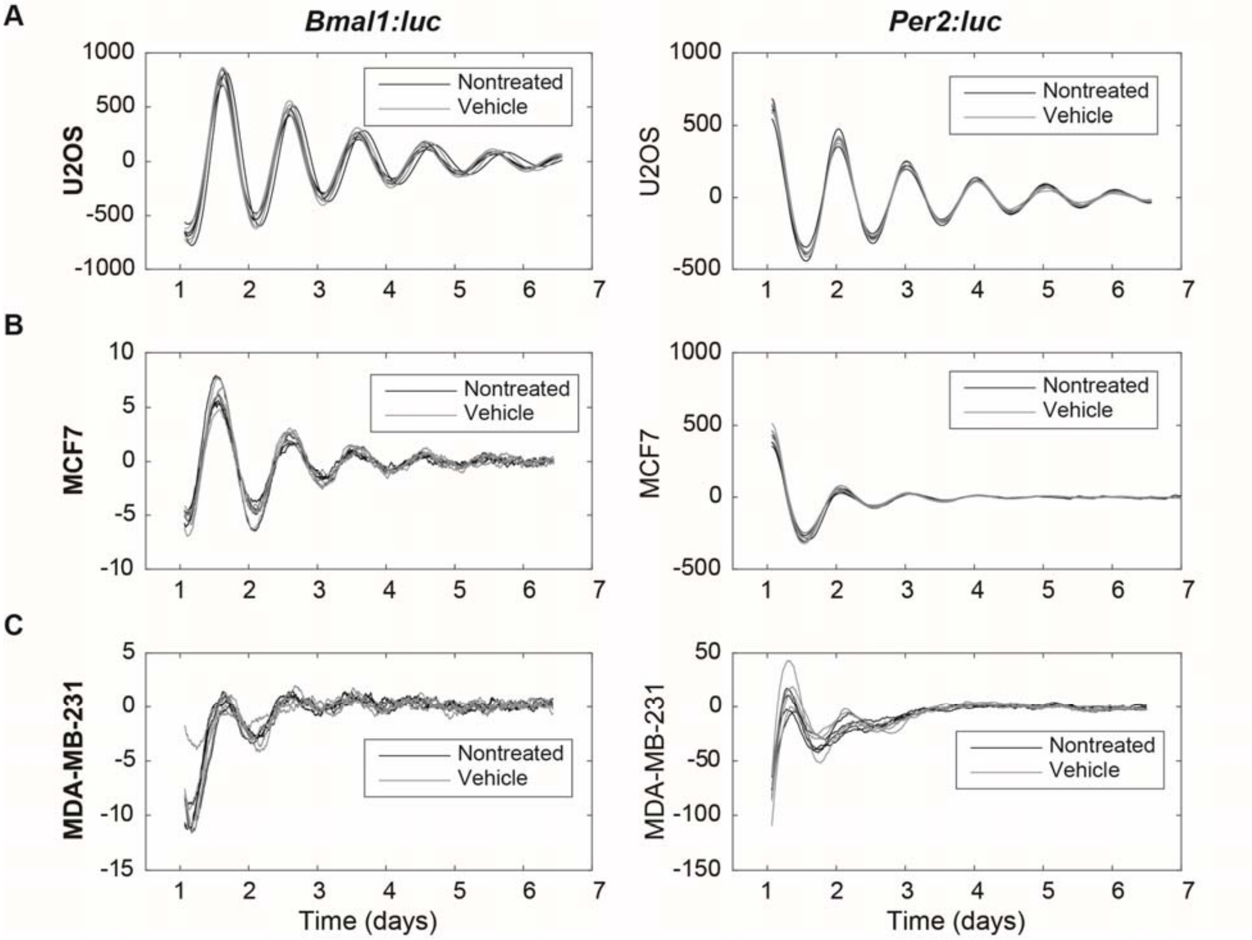


**Fig. S6. Vehicle treatment has no effect on circadian oscillations.** Shown are the smoothed and de-trended replicates for *Bmal1:luc* and *Per2:luc* reporters in (**A**) U2OS, (**B**) MCF7, and (**C**) MDA-MB-231 cells under non-treated (black) and vehicle-treated (gray) conditions. N=4 for each treatment for each cell line. Data were evaluated beginning from t=0.5 (day); each trace has been de-trended by subtracting the mean of a 24h sliding window and smoothed using the mean of a 3-h sliding window, making the de-trended date begin at t=1 (days).

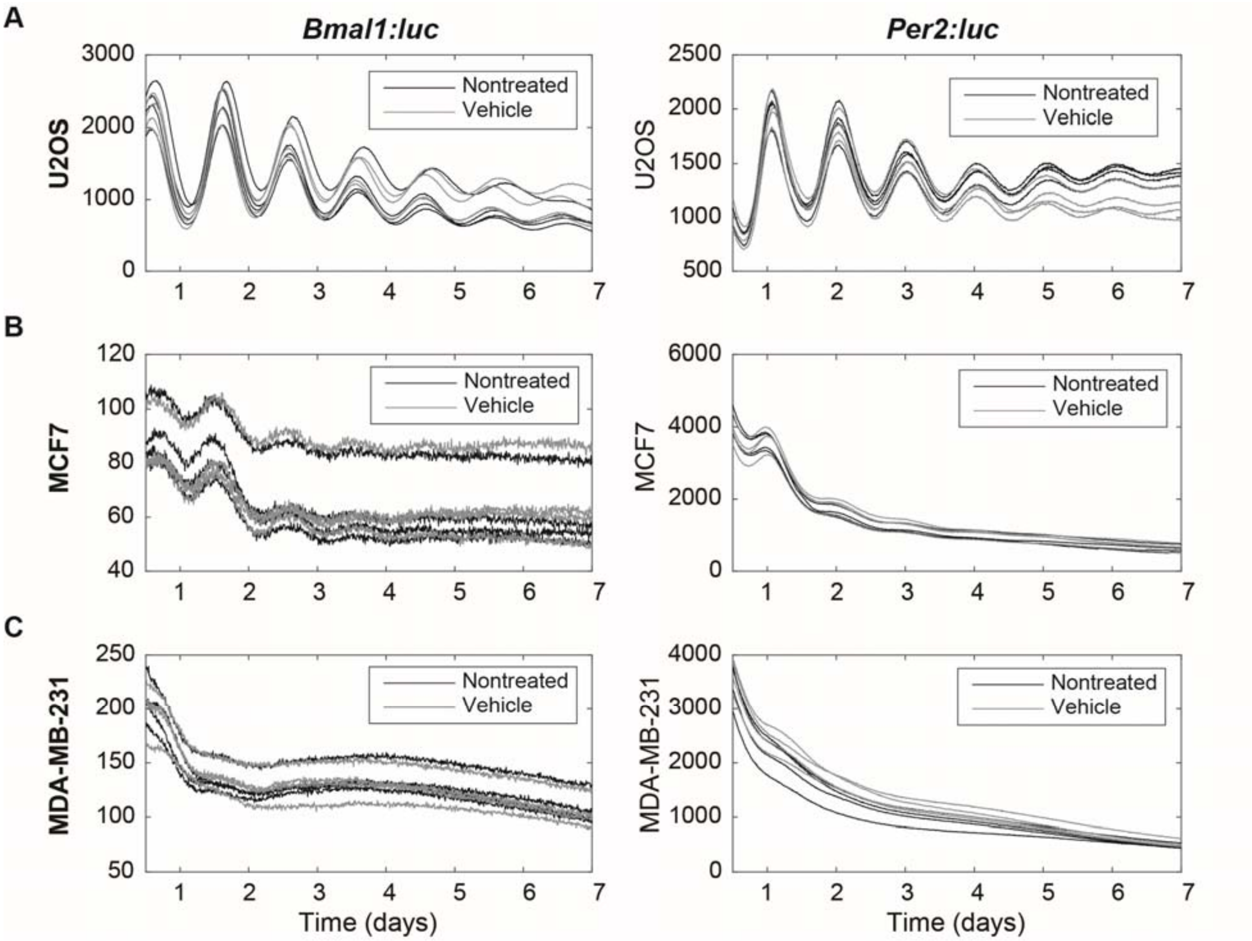


**Fig. S7. Vehicle treatment has no effect on circadian oscillations.** Shown are raw data replicates for *Bmal1:luc* and *Per2:luc* reporters in (**A**) U2OS, (**B**) MCF7, and (**C**) MDA-MB-231 cells under non-treated (black) and vehicle-treated (gray) conditions. N=4 for each treatment for each cell line.

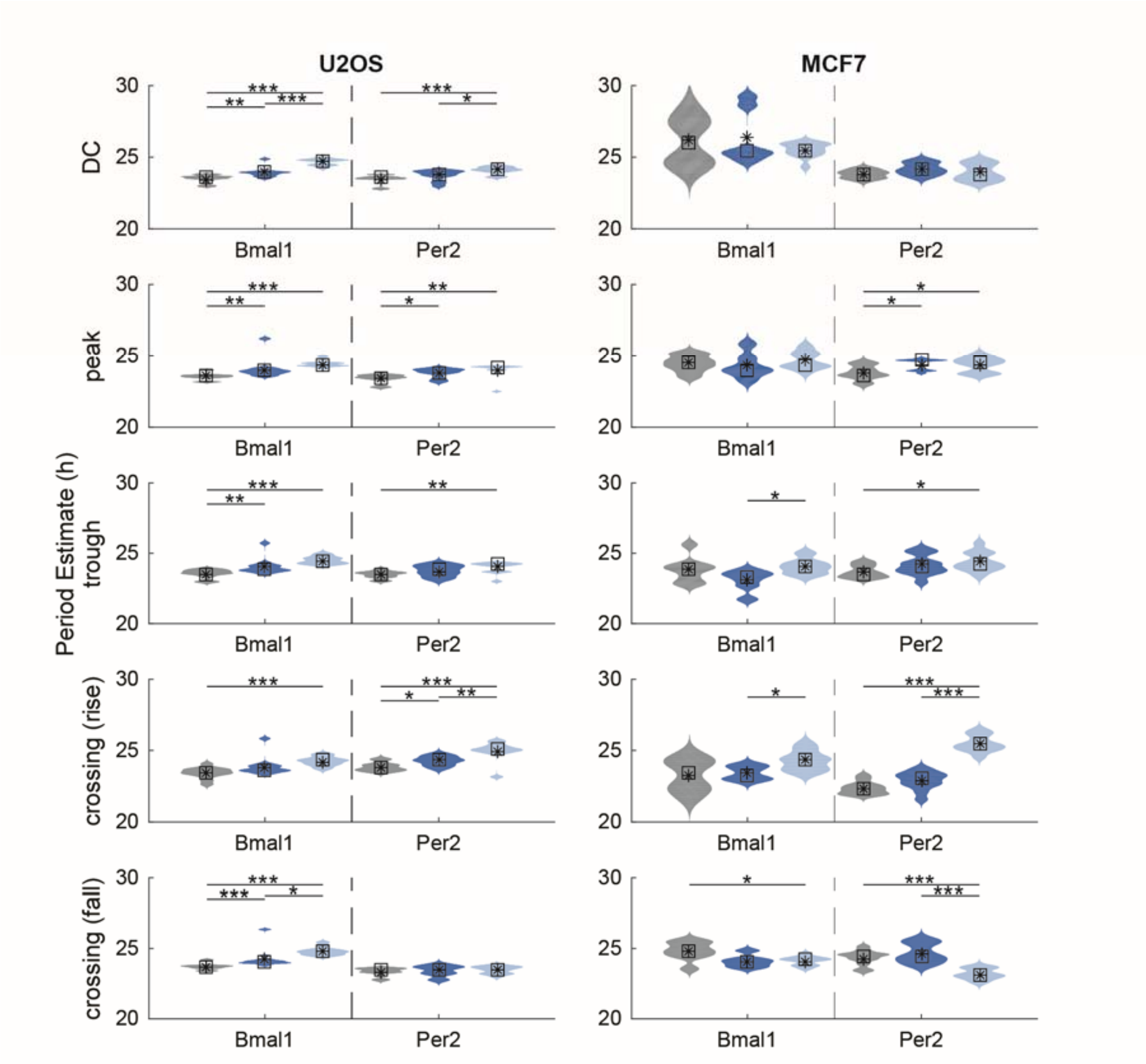


**Fig S8. Multiple period-estimation methods reveal similar trends for U2OS *Bmal1:luc* and *Per2:luc* and MCF7 *Per2:Luc* recordings, but not MCF7 *Bmal1:Luc*.** Shown are the distributions of period estimates for each recording (N=12 for each condition for each reporter and cell line). For each cell line (U2OS on left, MCF7 on right) and each reporter (*Bmal1* on left, *Per2* on right), we show the period distribution, color-coded by treatment (gray = vehicle, blue= 5 μM nobiletin, light blue = 50 μM nobiletin). Each row shows results from a different method (DC=damped cosine-fitting; the remaining method names indicate the phase marker used to estimate the period, e.g. "peak" indicates the difference in time between the peak of each cycle is used). Randomization tests for difference in means were performed to determine if treatment led to statistically significant differences. P-values were corrected according to Bonferroni's method (* $p < 0.05$, ** $p < 0.01$, *** $p < 0.001$). For U2OS (both reporters) and MCF7 *Per2:luc*, the period estimates of 50 μM nobiletin treated recordings are either longer than or not statistically significantly different from those treated with vehicle or 5 μM. For *MCF7:Bmal1*, however, two methods (trough-to-trough estimates and mean-crossing while levels are rising) indicate that the 50 μM nobiletin treatment leads to shorter periods and one (mean-crossing while levels are falling) indicates that it leads to longer periods.

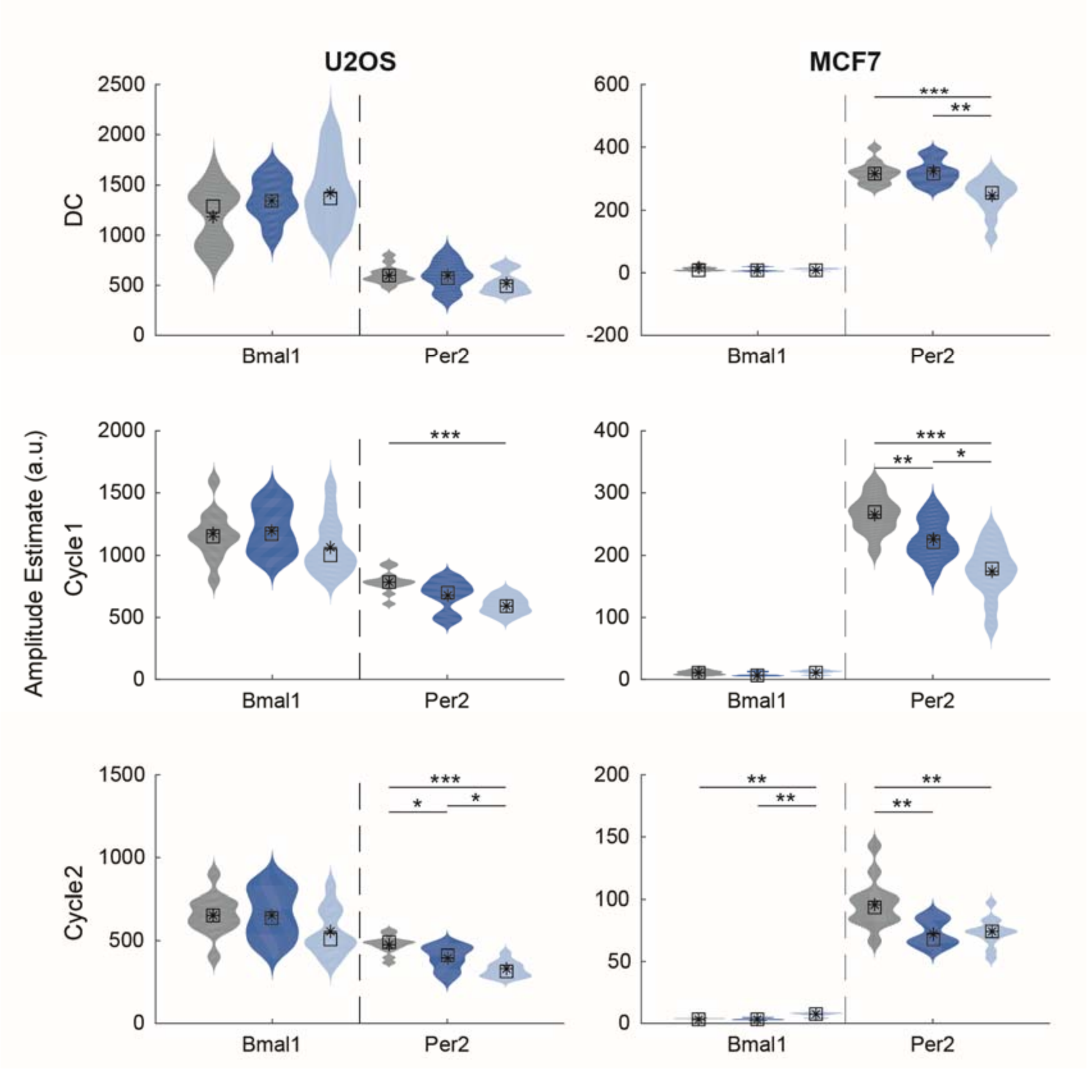


**Fig S9**. **Multiple amplitude-estimation methods reveal few trends for *Bmal1:Luc*, the same trends for U2OS *Per2:luc*, and conflicting trends for MCF7 *Per2:Luc*.** Shown are the distributions of amplitude estimates for each recording (N=12 for each condition for each cell line). For each cell line (U2OS on left, MCF7 on right) and each reporter (*Bmal1* on left, *Per2* on right), we show the amplitude distribution, color-coded by treatment (gray = vehicle, blue = 5 μM nobiletin, light blue = 50 μM nobiletin). Each row shows results from a different method (DC=damped cosine-fitting; Cycle 1= sum of magnitudes of first peak and first trough; Cycle 2 = sum of magnitudes of second peak and second trough). Randomization tests for difference in means were performed to determine if treatment led to statistically significant differences. P-values were corrected according to Bonferroni's method (* $p < 0.05$, ** $p < 0.01$, *** $p < 0.001$). For U2OS *Bmal1:luc*, there are no trends. For U2OS *Per2:luc*, 50 μM treatment led to a smaller amplitude. For MCF7 *Bmal1:luc*, only the amplitude of the second cycle is different across treatments, with a 50 μM treatment leading to a higher amplitude. For MCF7 *Per2:luc*, 50 μM treatment led to a smaller amplitude.

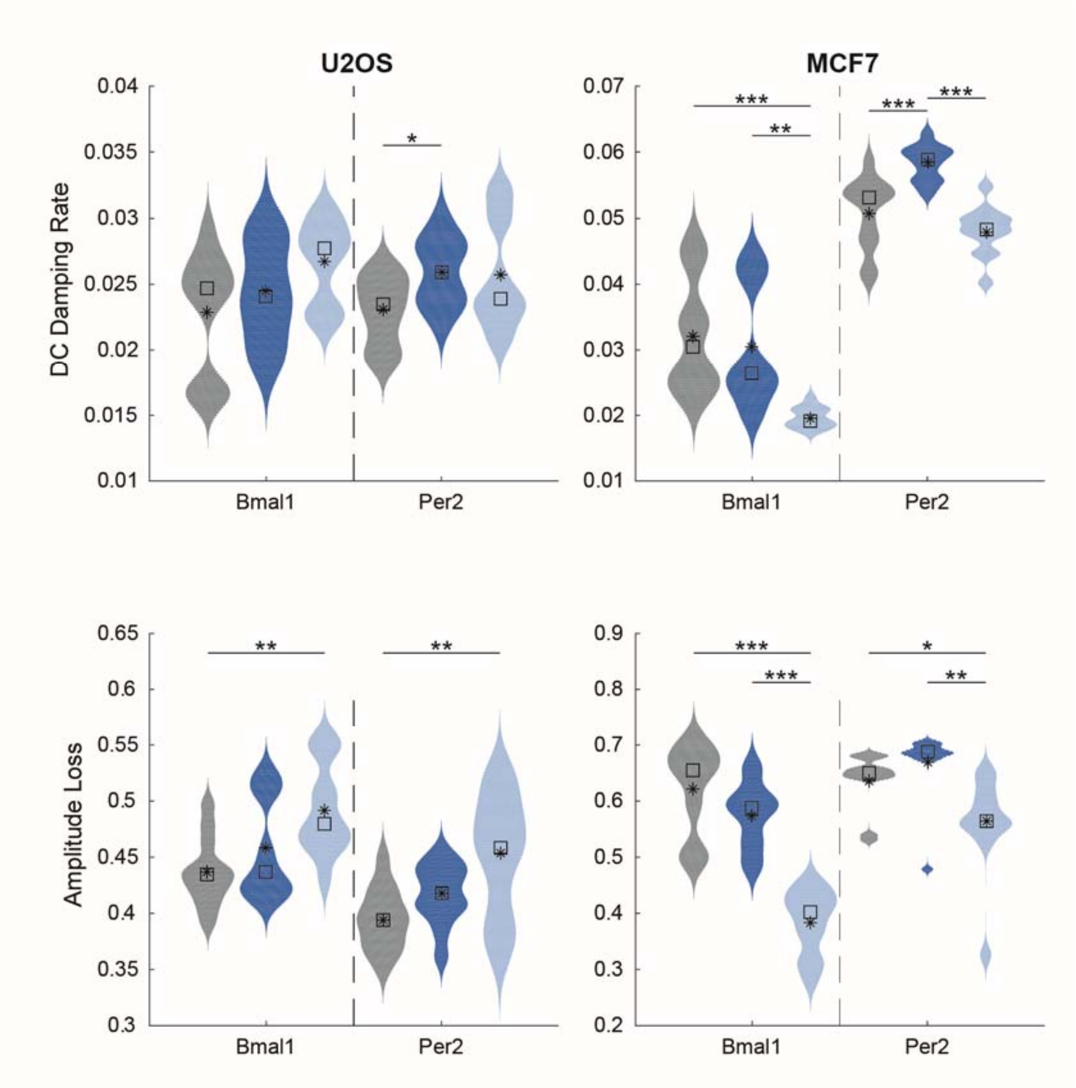


**Fig S10**. **Multiple damping-estimation methods reveal moderate trends for both U2OS *Bmal1:luc* and for MCF7.** Shown are the distributions of damping estimates for each recording (N=12 for each condition for each cell line). For each cell line (U2OS on left, MCF7 on right) and each reporter (*Bmal1* on left, *Per2* on right), we show the amplitude distribution, color-coded by treatment (gray = vehicle, blue = 5 μM nobiletin, light blue = 50 μM nobiletin). Each row shows results from a different method (DC Damping rate=damped cosine-fitting; Amplitude Loss = 1 – Cycle2Amplitude/Cycle1Amplitude). Randomization tests for difference in means were performed to determine if treatment led to statistically significant differences. P-values were corrected according to Bonferroni's method (* $p < 0.05$, ** $p < 0.01$, *** $p < 0.001$). For U2OS *Bmal1:luc* and *Per2:luc,* 50 μM nobiletin treatment leads to increased amplitude loss. For MCF7, 50 μM nobiletin treatment reduces damping (as determined both by cosine-fitting and by measuring amplitude loss from cycle 1 to cycle 2).

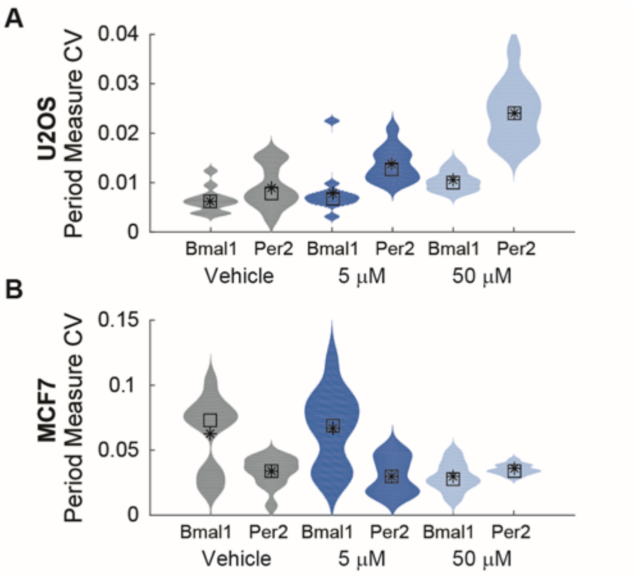


**Fig S11**. **Agreement of period-estimation methods is consistent for U2OS and and mildly inconsistent for MCF7.** For a given reporter and treatment, each of 12 replicates had its period estimated by 5 methods (damped-sine fitting, peak-to-peak, trough-to-trough, mean-crossing-to-mean-crossing on the rise, mean-crossing-to-mean-crossing on the fall). The coefficient of variation (CV) across the 5 methods is computed. Shown are the distributions of CVs for (**A**) U2OS and (**B**) MCF7 *Bmal1:luc* and *Per2:luc* recordings. Color indicates treatment (gray = vehicle, blue = 5 μM nobiletin, light blue = 50 μM nobiletin). For U2OS, the CV of period estimation is less than 4%. For MCF7 *Bmal1:luc*, vehicle treatment and 5 μM nobiletin period estimates vary more (the standard deviation is 10% of the mean).

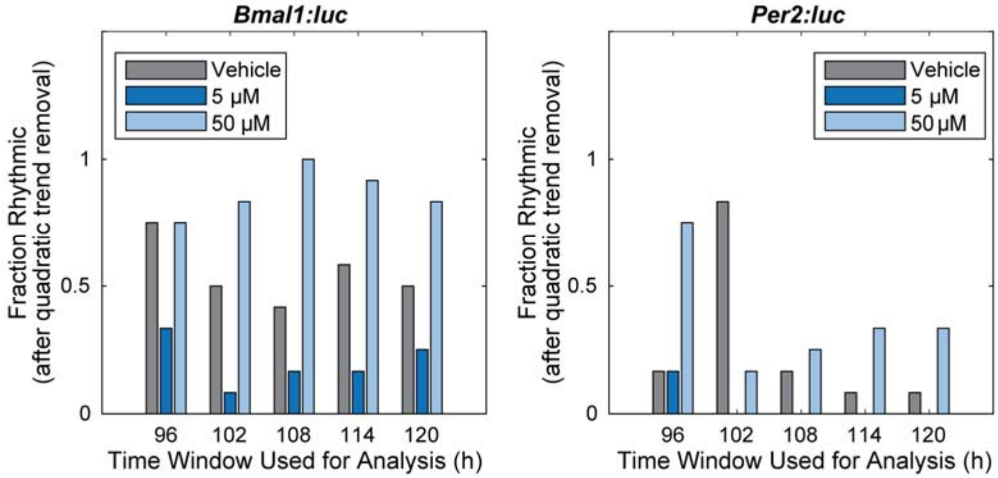


**Fig. S12**. **Nobiletin alters the rhythmicity of MDA-MB-231 cells.** Shown are the fractions of recordings classified as rhythmic ($p < 0.05$ on FFT-based test), after removing a quadratic trend, using time-series with increasing end times. The fraction scoring as rhythmic depends on the extent of the time-series included in the analysis, but cells treated with 50 µM nobiletin consistently scored as rhythmic more frequently than those treated with either vehicle or 5 µM. The only exception is for *Bmal1:luc* recordings with the first 96 hours analyzed.

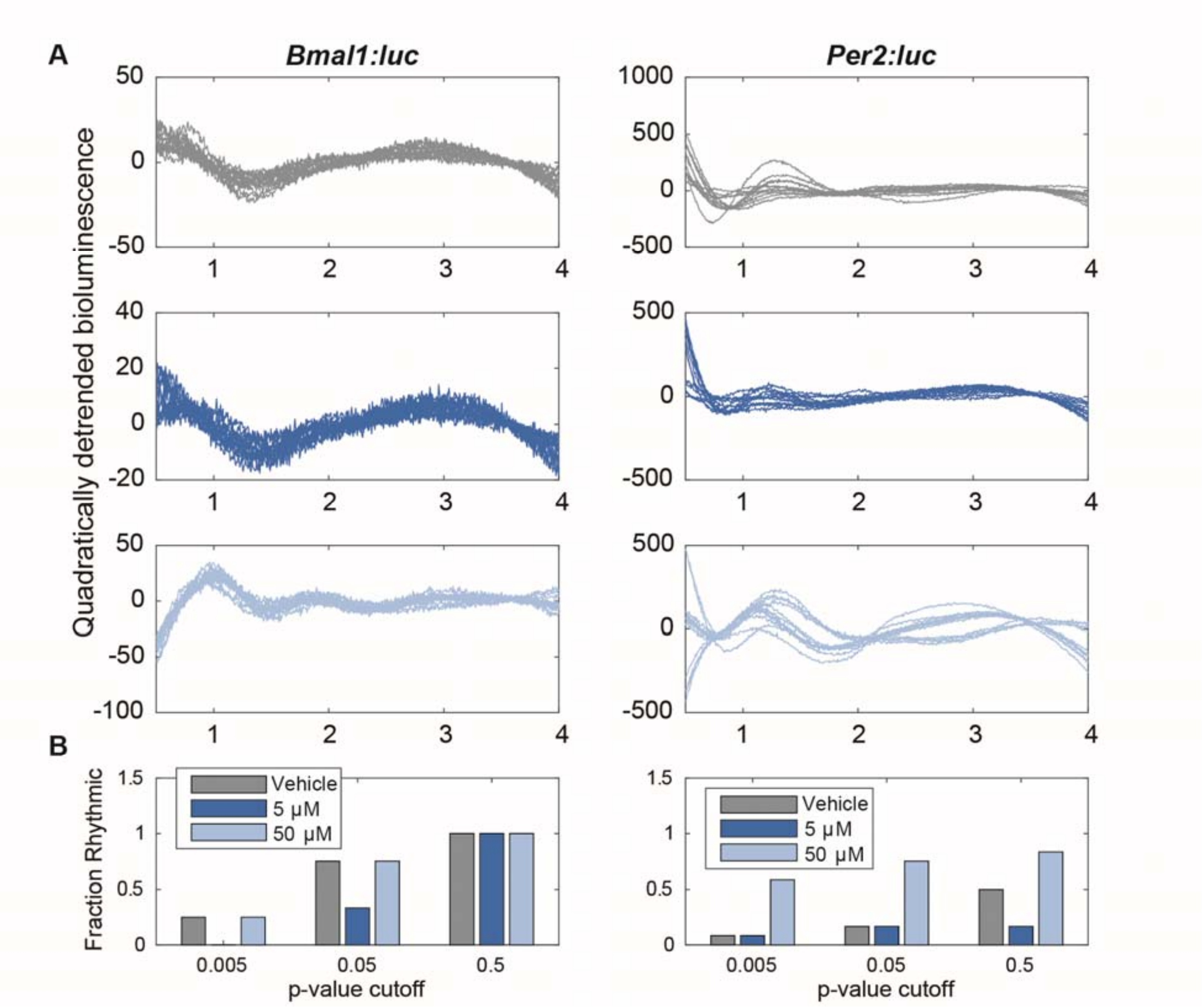


**Fig. S13**. **Removing a quadratic trend from the first four days shows that nobiletin alters the rhythmicity of MDA-MB-231 cells. (A)** Shown are luminometry recordings after removing a quadratic trend from the first 96 hours of data. Subfigure placement and color and indicate treatment (top/gray = vehicle, middle/blue = 5 μM nobiletin, bottom/light blue = 50 μM nobiletin). **(B)** Shown are the fractions of recordings classified as rhythmic by an FFT-based test, using increasing p-value cut-offs. For *Bmal1:luc*, both vehicle-treated and 50 μM nobiletin-treated cells are scored as rhythmic, but visual inspection demonstrates that the rhythm is most prominent for recordings from the 50 μM nobiletin treatment. For *Per2:luc*, visual inspection and the fraction scored as rhythmic both indicate that rhythms are present in cells treated with 50 μM nobiletin.

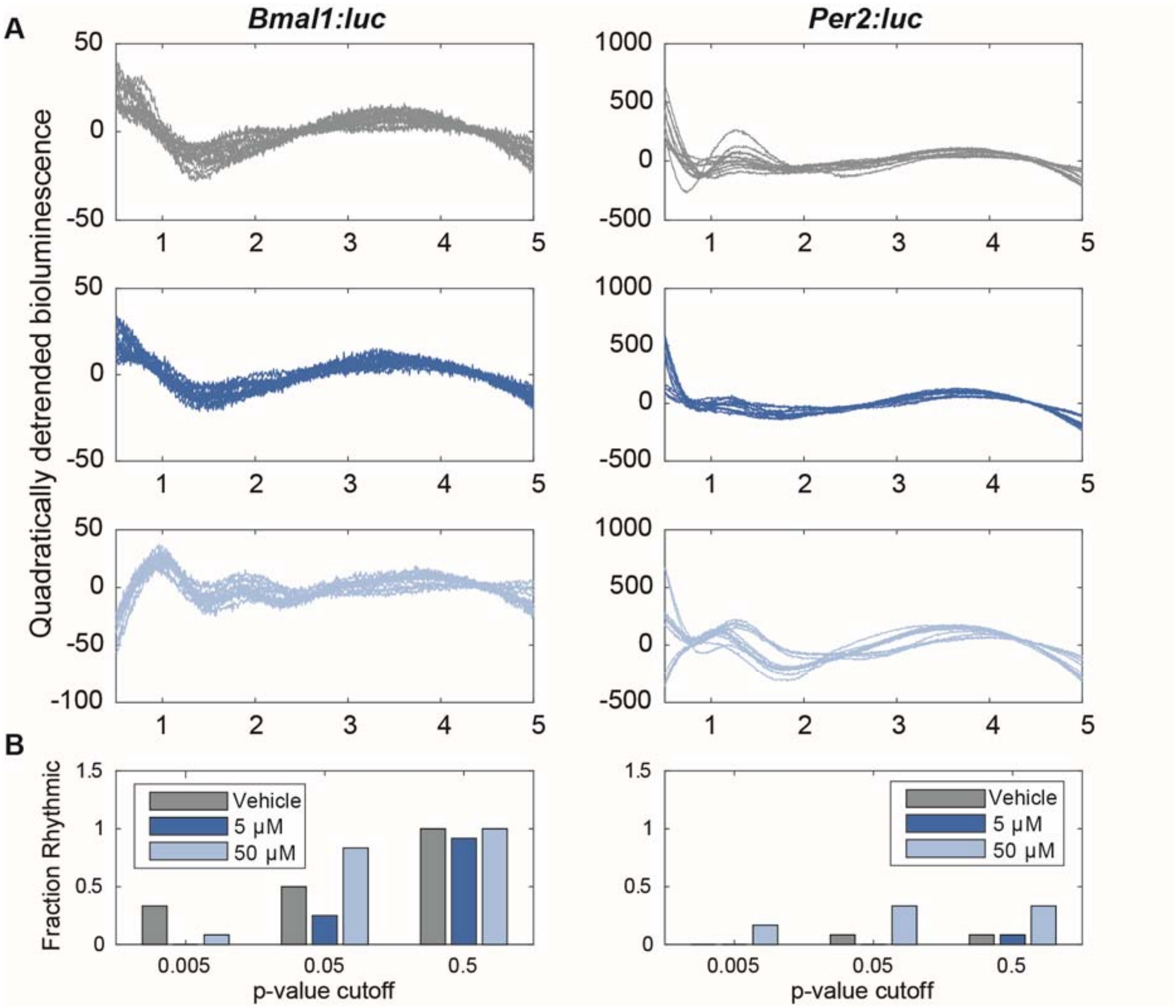


**Fig. S14**. **Removing a quadratic trend from the first five days shows that nobiletin alters the rhythmicity of MDA-MB-231 cells. (A)** Shown are luminometry recordings after removing a quadratic trend from the first 96 hours of data. Subfigure placement and color and indicate treatment (top/gray = vehicle, middle/blue = 5 μM nobiletin, bottom/light blue = 50 μM nobiletin). **(B)** Shown are fractions of recordings classified as rhythmic by an FFT-base test, using increasing p-value cut-offs. Visual inspection and the fraction scored as rhythmic (when p=0.05) both indicate that rhythms are present in cells treated with 50 μM nobiletin. For *Per2:luc*, this pattern is consistent, regardless of the size of the p-value. For *Bmal1:luc*, both vehicle-treated and 50 μM nobiletin-treated cells scored as rhythmic, but visual inspection demonstrates that the rhythm is most prominent for recordings from the 50 μM nobiletin treatment. For *Per2:luc*, visual inspection and the fraction scored as rhythmic both indicate that rhythms are present in cells treated with 50 μM nobiletin.

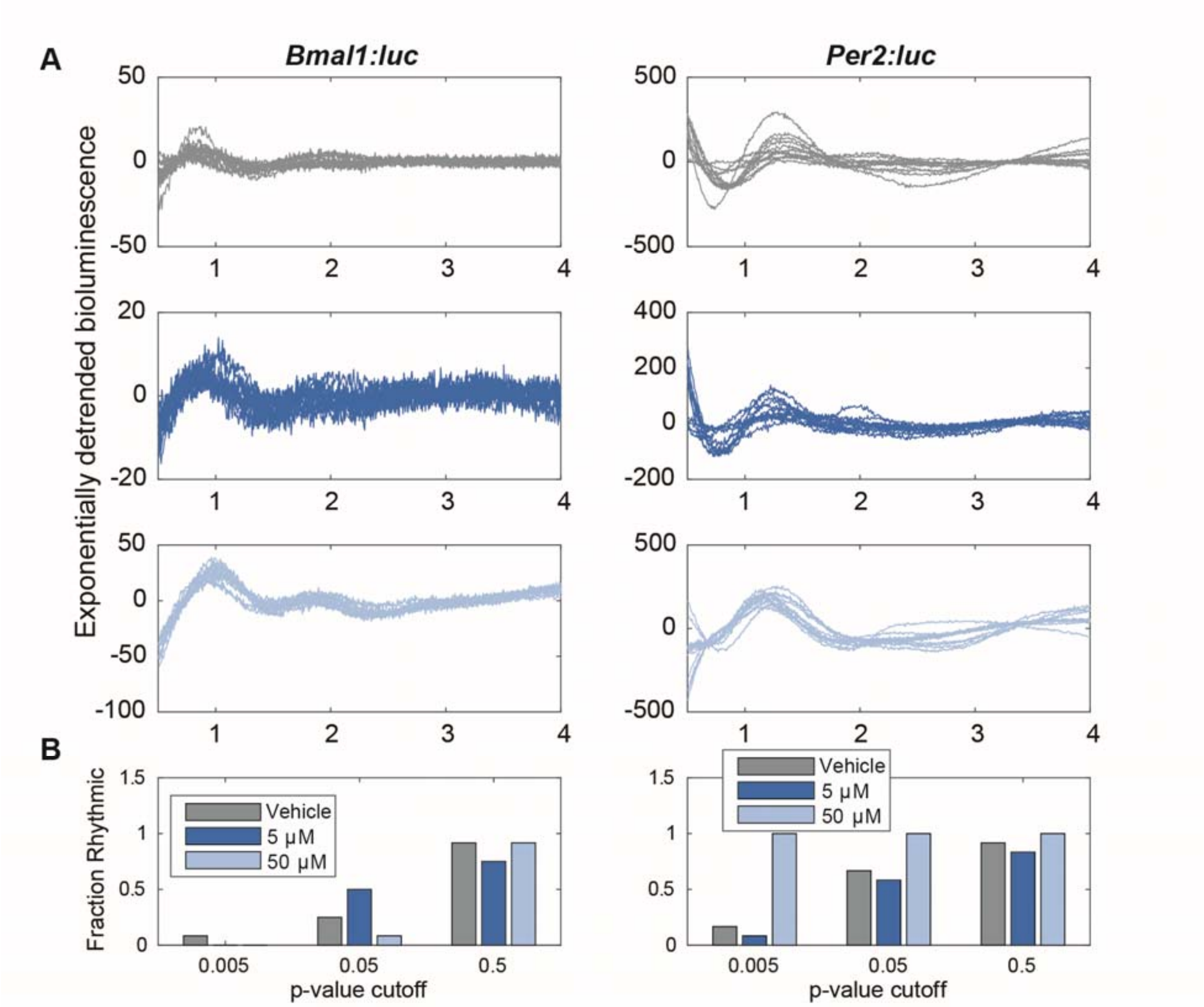


**Fig. S15**. **Removing an exponential trend from the first four days shows that nobiletin alters the rhythmicity of MDA-MB-231 cells. (A)** Shown are luminometry recordings after removing an exponential trend from the first 96 hours of data. Subfigure placement and color and indicate treatment (top/gray = vehicle, middle/blue = 5 μM nobiletin, bottom/light blue = 50 μM nobiletin). **(B)** Shown are fractions of recordings classified as rhythmic by an FFT-base test, using increasing p-value cut-offs. For *Bmal1:luc*, both vehicle- and 50 μM nobiletin-treated cells scored as rhythmic, but visual inspection demonstrates that the rhythm is most prominent for recordings from the 50 μM nobiletin treatment. For *Per2:luc*, visual inspection and the fraction scored as rhythmic both indicate that rhythms are present in cells treated with 50 μM nobiletin.

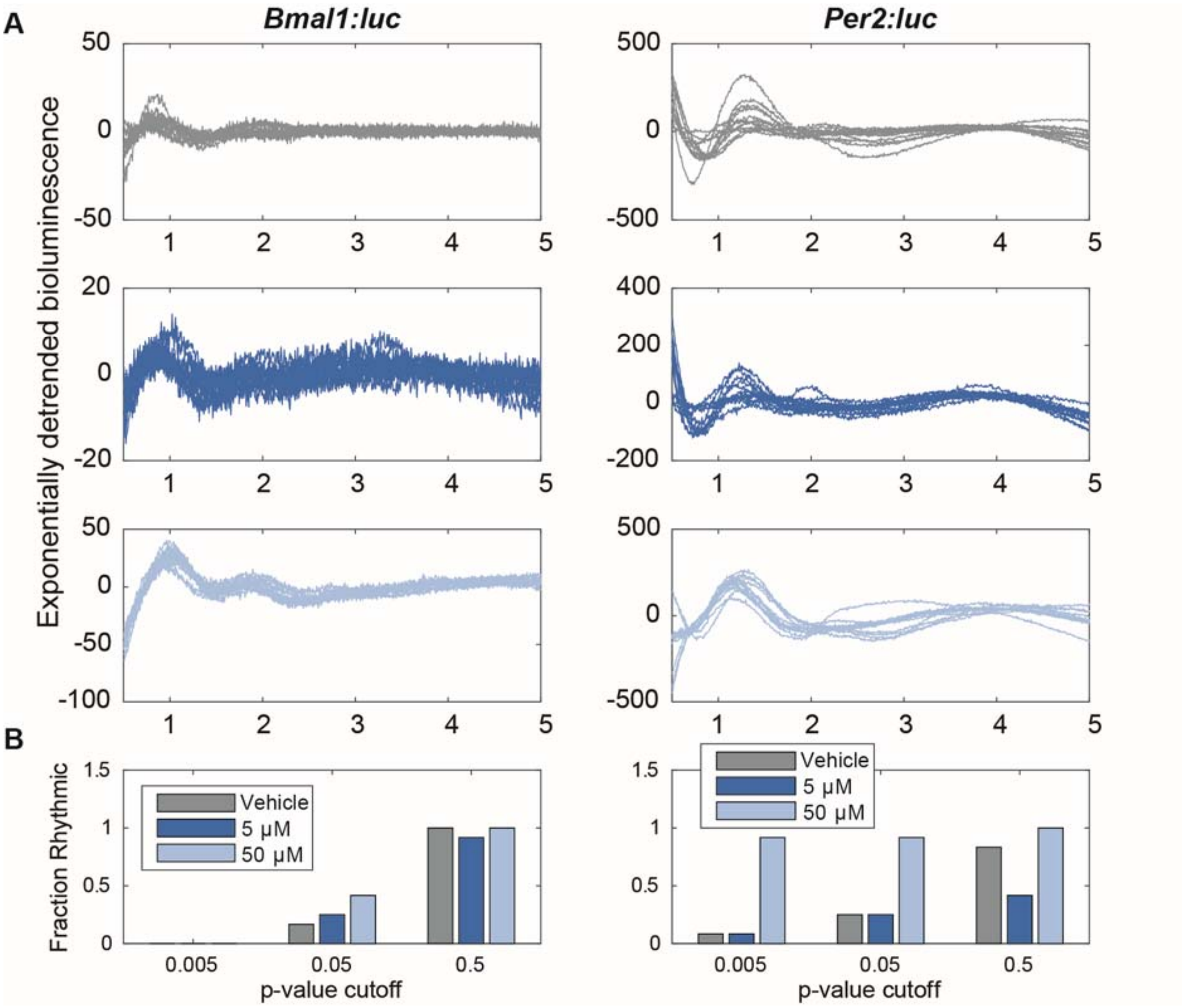


**Fig. S16**. **Removing an exponential trend from the first five days shows that nobiletin alters the rhythmicity of MDA-MB-231 cells. (A)** Shown are luminometry recordings after removing an exponential trend from the first 96 hours of data. Subfigure placement and color and indicate treatment (top/gray = vehicle, middle/blue = 5 μM nobiletin, bottom/light blue = 50 μM nobiletin). **(B)** Shown are the fractions of recordings classified as rhythmic by and FFT-base test, using increasing p-value cut-offs. Visual inspection and the fraction scored as rhythmic (when p=0.05) both indicate that rhythms are present in cells treated with 50 μM nobiletin. For *Per2:luc*, this pattern is consistent, regardless of the size of the p-value.

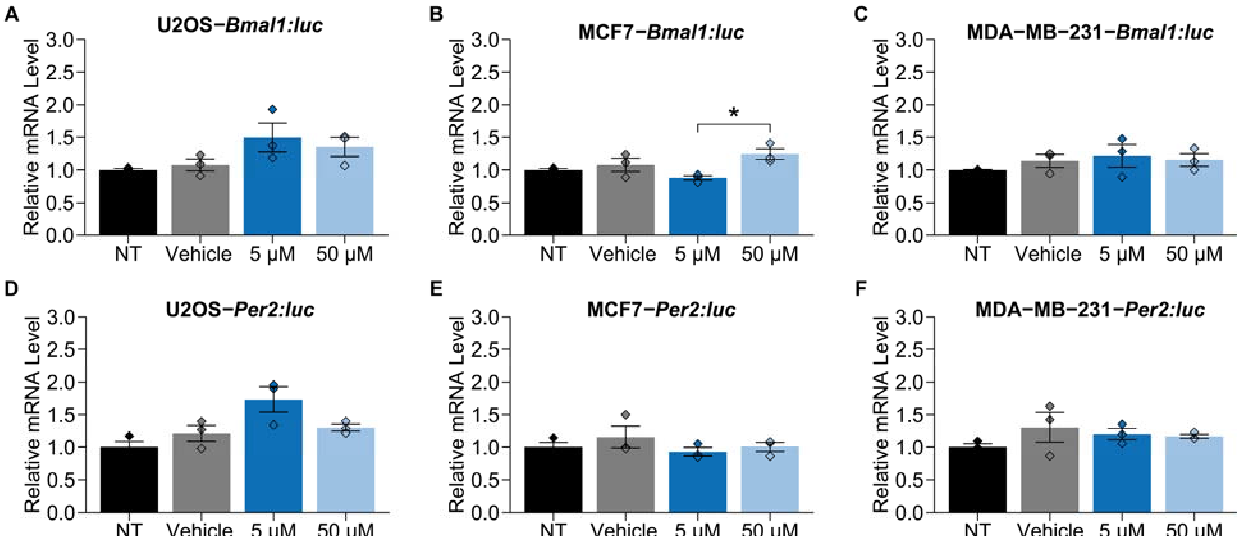


**Fig. S17**. **Nobiletin does not increase *Bmal1* or *Per2* mRNA levels in U2OS, MCF7, or MDA-MB-231 cells.** mRNA levels were quantified using RT-qPCR. No significant differences were observed in nobiletin treated samples compared to vehicle treated samples in any of the cell lines tested. Each treatment contained three biological replicates, with three technical replicates each. Error bars represent standard error of the mean (SEM). Statistical significance was evaluated via two tailed t.test in R ggpubr library under equal variance, and 0.95 confidence interval (*p < 0.05). NT = non-treated; Vehicle = 0.2% DMSO.

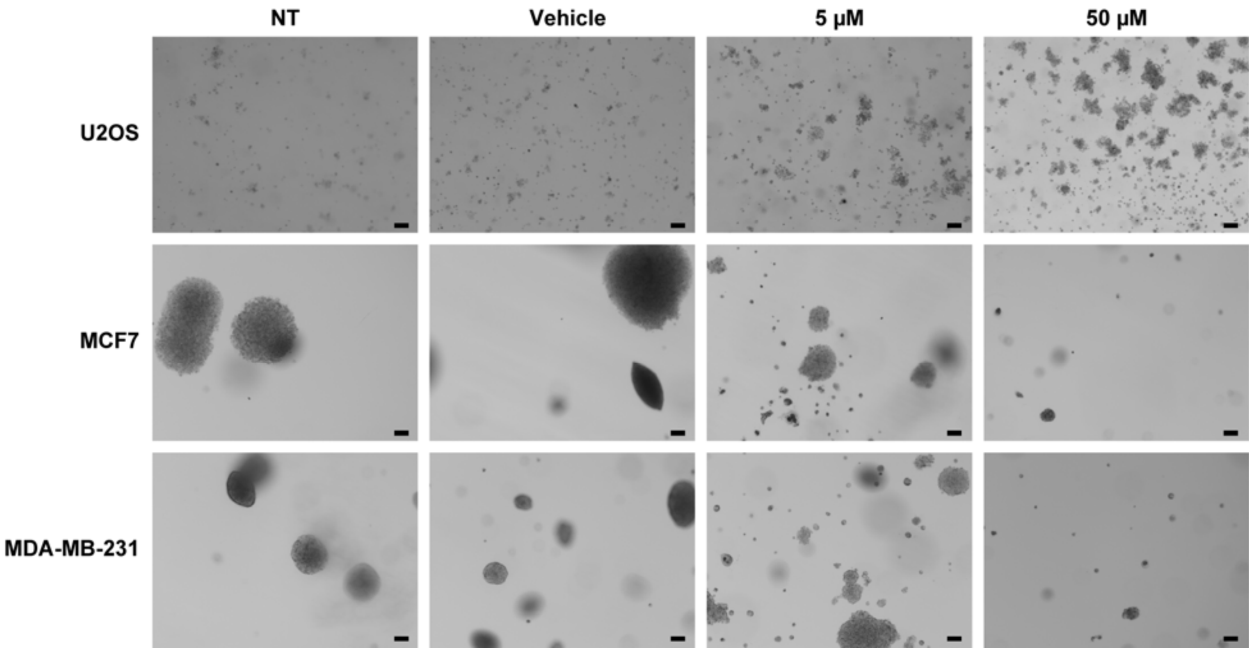


**Fig. S18**. **Representative images from colony formation assay.** A single image for each treatment and cell line is shown. Scale bars are 106 μm.
